# Transcriptomic analysis identifies synapse-enriched lncRNAs required for excitatory synapse development and fear memory

**DOI:** 10.1101/2023.07.14.549055

**Authors:** Sarbani Samaddar, Balakumar Srinivasan, Kamakshi Garg, Nandu Raj, Sania Sultana, Utsav Mukherjee, Dipanjana Banerjee, Wei-Siang Liau, Dasaradhi Palakodeti, Timothy W. Bredy, Sourav Banerjee

## Abstract

Regulatory functions of lncRNAs in neurons have been majorly limited to the nucleus. The identity of synaptic lncRNAs and their functional roles associated with synapse development and plasticity are poorly understood. Transcriptomic analysis from synaptoneurosomes identified 94 synapse-enriched lncRNAs from the mouse hippocampus. We find Pvt1 to be a specific regulator of excitatory, not inhibitory, synapse development. Synapse-specific loss of Pvt1 regulates synaptic activity and the downregulation of mRNAs pertinent to synapse development. We also report a synapse-centric role for an uncharacterized lncRNA; 2410006H16Rik (SynLAMP); in translation and memory formation. SynLAMP demonstrates enhanced localization to the synapse following fear conditioning and regulates dendritic translation by sequestering the translation repressor FUS; acting as a molecular decoy. Synapse-specific knockdown of SynLAMP inhibits the localised, activity-dependent translation of CamK2a, a FUS target. SynLAMP RNAi occludes fear-memory formation. Comprehensively, our study highlights that *de novo* activity of lncRNAs are involved in diverse synaptic functions.

## Introduction

Within neurons, functional autonomy of subcellular compartments such as synapses rely on the *in-situ* availability of proteins to meet the requirements for synaptic development and plasticity. Localization of RNAs to the synapse enables neurons to rapidly fulfill such on-site demands for stimulus-specific translation in a spatiotemporal manner (Holt and Schuman, 2013; Martin and Ephrussi, 2009; Sutton and Schuman, 2006). Local translation from synaptically available coding transcripts in response to stimulus are physiologically critical for learning and memory (Ashraf et al., 2006; Chen et al., 2022; Conde-Dusman et al., 2021; Wang et al., 2009). A number of regulatory elements are therefore present *in situ* at the synapse in order to monitor the diversity and dynamicity of the synaptic proteome.Transcriptomic analysis of the hippocampus neuropil has identified ∼2550 mRNAs in excitatory synapses (Cajigas et al., 2012). Regulatory RNAs, predominantly comprised of non-coding transcripts, such miRNAs (Kye et al., 2007; Siegel et al., 2009), circular RNAs (Piwecka et al., 2017; Rybak-Wolf et al., 2015; You et al., 2015). Diverse isoforms of 3’ untranslated region (3’UTR) (Tushev et al., 2018) have also been identified from the synapto-dendritic compartment. These regulatory RNAs, in association with RNA binding proteins (RBPs), determine the transport, stability, and translation of mRNAs associated with neuronal plasticity (Darnell, 2013; Holt and Schuman, 2013) and memory (Kandel, 2001; Sutton and Schuman, 2006). However, no such detailed characterization exists for long noncoding RNAs (lncRNAs).

*In situ* hybridization data from the Allen Brain Atlas have identified 849 lncRNAs that are expressed either ubiquitously or in specific neuroanatomical regions of the brain (Mercer et al., 2008). This, together with findings from GENCODE (Harrow et al., 2012) and NONCODE (Bu et al., 2012; Xie et al., 2014 dismissed the preexisting notion that lncRNAs are simply by-products of transcription (Briggs et al., 2015; Samaddar and Banerjee, 2021). From a functional perspective, our understanding of brain-enriched lncRNAs primarily stems from nuclear lncRNAs. Almost all the neuronal lncRNAs reported to date are restricted to the nucleus, (Clemson et al., 2009; Khalil et al., 2009; Mondal et al., 2010; Tsai et al., 2010; Wei et al., 2022; Zhao et al., 2019) with a few exceptions; the lncRNAs AK077867 (Mercer et al., 2008), and ADEPTR (Grinman et al., 2021), SLAMR (Espadas et al., 2024) and select isoforms of Gas5 (Liau et al., 2023). The number and nature of cytoplasmic lncRNAs localized at the synapse remain undocumented thus far. Here, we performed a detailed transcriptomic analysis of the synaptic compartment of the adult mouse hippocampus to identify the full repertoire of lncRNAs localized at the synapse. We found 1031 lncRNAs, including a subset of 94 lncRNAs that were significantly enriched in the synaptic compartment.

Amongst the synaptically enriched lncRNAs, we focused on the characterization of a few synaptically abundant lncRNAs to define their role in functional synapse development and synaptic plasticity. We have investigated the somato-dendritic distribution of lncRNAs and the dynamics of lncRNA-RBP interaction in contextual fear conditioning. Furthermore, we have explored the involvement of these lncRNAs in dendritic protein synthesis and memory.

## Results

### Identification of synapse-enriched long non-coding RNAs in the hippocampus

To identify the full repertoire of hippocampal synapse-enriched lncRNAs, we sequenced total RNA extracted from purified synaptoneurosomes. As synaptoneurosomes are highly enriched for synaptic membranes that preserve components of the translation machinery and regulatory RNAs including non-coding transcripts (Rao and Steward, 1991; Schratt et al., 2004; Siegel et al., 2009), we reasoned that they represent an appropriate source of synapse-enriched lncRNAs. Synaptoneurosomes were purified from the mouse hippocampus (8 week old) using a Ficoll gradient (Figure 1A), with the purity of this biochemical fraction being assessed by the enrichment of synaptic proteins PSD95 and Synaptophysin, along with the depletion of nuclear protein Histone H1 and astrocytic protein GFAP (Figure 1B). Three independent sequencing runs were performed using total RNA purified from synaptoneurosomes and cell soma.GENCODE (M18) was used to identify lncRNAs from mapped reads.

**Figure 1:**
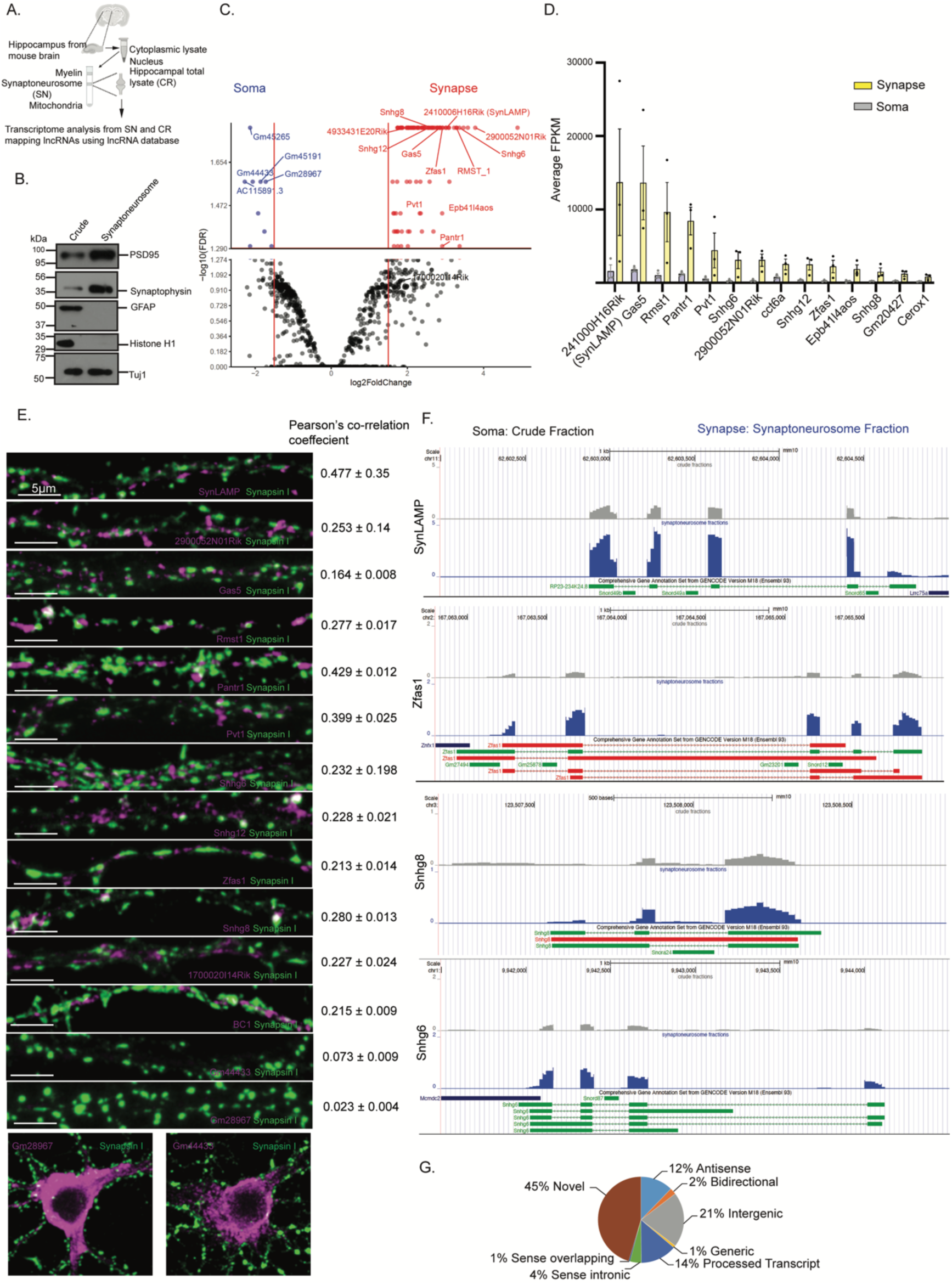
Identification and localization of lncRNAs at the hippocampal synapse. (**A**) Schematic representation of synaptoneurosome preparation (SN) from adult mouse hippocampus and identification of synapse-enriched lncRNAs by RNASeq analysis from 3 independent experiments. (**B**) Evaluation of SN quality by western blot. (**C**) Volcano plot showing lncRNAs identified from hippocampal neurons. Highlights indicate lncRNAs significantly (FDR<0.05) enriched at the synapse (red) and soma (blue). (**D**) Expression of highly abundant and evolutionarily conserved lncRNAs in soma (grey) and synapse (yellow). (**E**) FISH showing localization of lncRNAs at the synapse and soma (magenta puncta). Co-localization (white puncta) showing synaptic localization of lncRNAs. Pearson’s correlation co-efficient and scale as indicated. n= 10-15 per lncRNA from 3 independent experiments. Also see Figure S1 **(F)** Custom-tracks from UCSC Genome Browser showing expression of indicated lncRNAs in soma and synaptoneurosome. Synaptic lncRNAs lack snoRNA signatures. (**G**) Categorization of lncRNAs identified from SN shown in pie chart with indicated percentage of novel and diverse types of annotated lncRNAs. The annotated lncRNAs are classified based on their genomic location. Also see Figure S2.

We have identified 563 annotated (55% of total transcripts) and 468 (45% of total transcripts) novel lncRNAs that are expressed in the hippocampus. An FPKM cut off higher than 10 was set to include lncRNAs that showed appreciable expression levels across all three independent transcriptomics experiments (Table S1). Among the annotated lncRNAs, 94 were significantly enriched at the synapse (FDR <0.05) (Figure 1C and 1D). Our study also detected 45 soma-enriched lncRNAs, among which, 9 of them showed significant enrichment (FDR <0.05). Identification of previously established synaptically enriched coding transcripts in our dataset further bolstered the accuracy of the transcriptomic analysis from synaptoneurosomes (Figure S2A). The synaptic localization of a subset of synapse-enriched lncRNAs was verified by *in situ* hybridization (Figures 1E and S1). A caveat of lncRNAs is that many act as hosts to snoRNAs that remain embedded within their sequences. To verify that the observed synaptically-enriched lncRNAs are not merely nuclear-origin transcripts that are transported to the synapse, we verified that the synapse-enriched isoforms of the lncRNAs are devoid of embedded snoRNAs (Figure 1F). This further corroborated our hypothesis that there exists a unique pool of lncRNAs within synapses.

A detailed analysis of all the lncRNAs is uploaded on the UCSC Genome Browser (http://genome.ucsc.edu/cgibin/hgTracks?db=mm10&hubUrl=https://cloud.rdm.uq.edu.au/Bags/Q5104/CR_SN.txt). Based on their genomic location with respect to protein-coding genes; lncRNAs are classified as antisense, intergenic, sense overlapping, sense intronic, bidirectional and processed transcripts (Dunham et al., 2012; Luo et al., 2020; Mercer et al., 2008, 2009). We have identified a significant proportion of intergenic (21%, 213), processed transcripts (14%, 141) and antisense (12%, 126) lncRNAs (GENCODE M18) (Figure 1G). We selected 2410006H16Rik (SynLAMP or <u>Syn</u>aptic<u>l</u>ncRNA <u>a</u>ssociated with <u>m</u>emory and <u>p</u>lasticity), Pantr1 and Pvt1 for further characterization based on their abundance in the synaptic compartment. There exists multiple spliced-isoforms of lncRNAs within neurons (Liau et. al. 2023); consistent with this observation, our transcriptomic analysis revealed, isoform specific synaptic distribution (Figure S2B).

### Functional analysis of synapse-enriched lncRNAs identifies Pvt1 as a regulator of dendrite and spine morphology

The structural alteration of dendritic spines correlates with the maturity and strength of excitatory synapses (Matsuzaki et al., 2004; Zito et al., 2004). To investigate the importance of lncRNAs in regulating dendritic arborization and spine structure, we have inhibited the functions of SynLAMP, Pantr1 and Pvt1 in hippocampal neurons by shRNA–mediated RNA interference (RNAi). Two distinct shRNAs were designed against each transcript and their efficacy determined using qRT-PCR (for Pantr1: 55.29± 14.61% decrease, p<0.005, for shRNA#1; 65.33 ± 13.07% decrease, p<0.0001, for shRNA#2; for Pvt1: 46.71 ± 12.58% decrease, p<0.0003, for shRNA#1, and 52.73 ± 11.03% decrease, p<0.0002, for shRNA#2; for SynLAMP: 44.94 ± 8.93 % decrease, p<0.0004, for shRNA#1, 69.37 ± 10.95% decrease, p<0.0001 for shRNA#2; Figure S3A-S3C). Plasmids expressing shRNA against indicated lncRNAs and EF1ɑ-mCherry were co-transfected with EGFP (CAG-EGFP) into primary neuronal cultures (DIV 3-4). Effective RNAi was determined by mCherry expression and dendritic arborization was calculated at DIV-21 based on EGFP fluorescence intensity. Pvt1 RNAi reduced dendritic arborization (distance from the soma: 70μm, 6.292±1.609; p<0.03; 80μm, 6.792±1.481, p<0.008; 90μm, 6.000±1.435, p<0.019; 100μm, 6.500±1.588, p<0.022); whereas SynLAMP and Pantr1 had no effect (Figure 2A - 2B). The loss of Pvt1 function also led to a decrease in spine density (spine numbers per μM of dendrite;Control RNAi: 1.032 ± 0.06, Pvt1 RNAi: 0.82 ± 0.05, p < 0.03) (Figure 2C - 2D).

**Figure 2:**
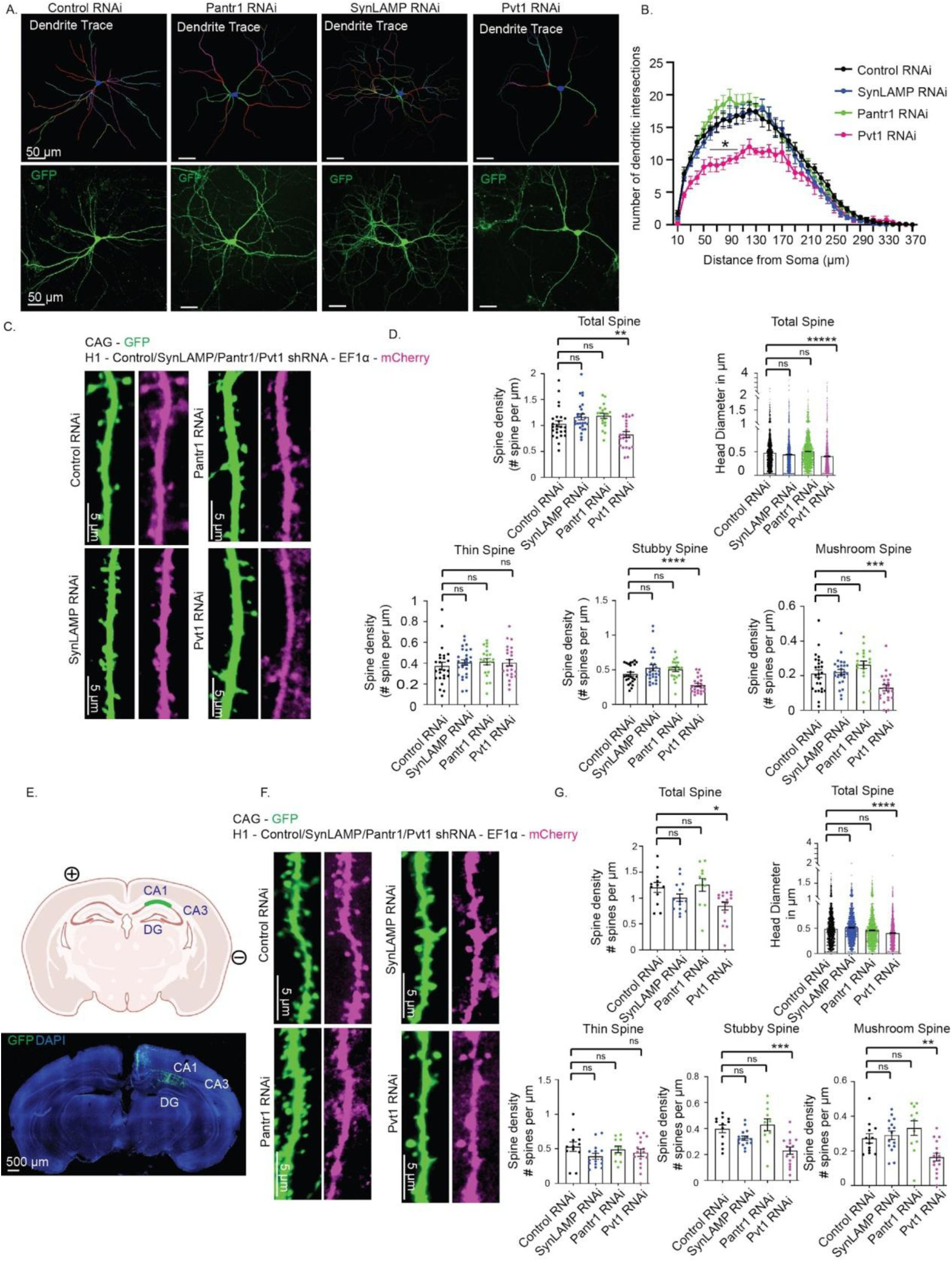
Pvt1 influences dendritic branching and spine growth. (**A**) Traces of reconstructed (upper panel) and confocal images (lower panel) of cultured neurons (DIV21) expressing non-targeting control shRNA or shRNA targeting Pantr1, SynLAMP and Pvt1 respectively. Scale as indicated. (**B**) Quantification of dendritic branching. n: control RNAi (12), Pantr1 RNAi (15), SynLAMP RNAi (20), Pvt1 RNAi (8) from 4-8 independent experiments in each condition, *p< 0.03. Multiple t-test determined by Holm-Sidak method. Data shown as mean ± SEM. (**C**) Images showing dendritic spines from cultured neurons expressing control shRNA or shRNA against Pantr1/SynLAMP/Pvt1 (as indicated). (D) Quantification of spine density, spine head diameter, and density of spine types. n = 1300 to 1900 spines from 8 to 12 neurons *in-vitro*, from 3 independent experiments *p<0.04, **p<0.03, ***p<0.008, ****p<0.002, *****p<0.0001, ns = not significant. One way Anova and Bonferroni correction. Data shown as mean ± SEM. (E) Plasmids harboring control shRNA or shRNA against Pantr1/SynLAMP/Pvt1 were electroporated into E17 embryos along with GFP. Brain slices were prepared at P28 for spine analysis from CA1 neurons. (**F**) Images showing dendritic spines of *in utero* electroporated CA1 neurons expressing indicated shRNA. (**G**) Quantification of spine density, spine head diameter, and spine types. n = 1350 to 1470 spines from 5 to 8 neurons *in vivo* from 3 independent experiments; *p<0.01, **p<0.02, ***p<0.0006, ****p<0.0001, ns = not significant. One way ANOVA and Bonferroni correction. Data shown as mean ± SEM. Also See Figure S3.

Knockdown of Pvt1 reduced the head diameter (Control RNAi: 0.47 ± 0.008 μM, Pvt1 RNAi: 0.40 ± 0.01 μM, p < 0.0001) of dendritic spines (Figure 2C - 2D). Analysis of spine types revealed that Pvt1 knockdown reduced the density of stubby (Control RNAi: 0.43 ± 0.02, Pvt1 RNAi: 0.28 ± 0.02, p < 0.002) and mushroom spines (Control RNAi: 0.21 ± 0.02, Pvt1 RNAi: 0.129 ± 0.01, p < 0.008) without affecting thin spine density. Although Pantr1 RNAi led to a miniscule increase in head diameter, the knockdown of Pantr1 and SynLAMP did not profoundly influence spine structure or type (Figure 2D). To complement our observations from *in vitro* studies, we also determined the spine structure and type from CA1 neurons in the hippocampus *in vivo* (P28) following *in utero* electroporation of SynLAMP, Pantr1 and Pvt1 shRNAs in CA1 neurons (E17) (Figure 2E). As in cultured neurons, knockdown of Pvt1 reduced the spine density (total spine density: Control RNAi: 1.2 ± 0.09, Pvt1 RNAi: 0.84 ± 0.07, p < 0.018; mushroom spine density: Control RNAi: 0.27 ± 0.02, Pvt1 RNAi: 0.16 ± 0.02, p < 0.02; and stubby spine density: Control RNAi: 0.398 ± 0.03, Pvt1 RNAi: 0.23 ± 0.02, p < 0.0006), but the density of the thin spines was not affected (Figure 2F - 2G). Pvt1 knockdown led to the reduction of head diameter (Control RNAi: 0.48 ± 0.007 μm, Pvt1 RNAi: 0.4 ± 0.006 μm, p < 0.0001) (Figure 2G). Consistent with our *in vitro* studies, knockdown of SynLAMP and Pantr1 had no profound effect on either spine structure or spine type *in vivo* (Figure 2F - 2G). These observations indicate that Pvt1 specifically regulates the morphology of dendritic spines that are sites for synapse formation.

### Pvt1 regulates excitatory synapse development

We next investigated the importance of SynLAMP, Pantr1 and Pvt1 in glutamatergic synapse development by measuring synapse density following their knockdown. Cultured hippocampal neurons (DIV 3 - 4) were transduced with lentivirus expressing each shRNA. Excitatory synapse density was measured from transduced glutamatergic neurons (DIV 21) on the basis of colocalization of Synapsin I and PSD95. The age of the neuron was commensurate with previous studies reporting a similar temporal window for synapse development (Fletcher et al., 1994; Paradis et al., 2007). Pvt1 knockdown led to reduced excitatory synapse density (Control RNAi: 1 ± 0.06, Pvt1 RNAi: 0.75 ± 0.07, p < 0.03), whereas the knockdown of Pantr1 and SynLAMP had no effect (Figure 3A-3B, S4A). To determine whether Pvt1 was involved in the regulation of both excitatory and inhibitory synapse development *in vivo*, Pvt1 RNAi was carried out in CA1 neurons *via in utero* electroporation (E17 embryos). Synapse density was measured (P28) on the basis of the colocalization of Synapsin I and PSD95 (for excitatory synapses) (Figure 3C-3D, S4B); and GAD65 and GABA (for inhibitory synapses) (Figure 3E-3G). We observed a significant reduction in excitatory synapse density (Control RNAi: 1 ± 0.08, Pvt1 RNAi 0.68 ± 0.03, p < 0.001) (Figure 3C - 3D). We did not observe any changes to the PSD95 and Synapsin I puncta density following the knockdown of Pvt1, Pantr1 and SynLAMP (Figure S4D-S4E). Pvt1 RNAi has no effect on inhibitory synapse density; either in the perisomatic regions or in the dendrites (Figure 3E – 3G, S4F). Together, these data indicate that Pvt1 is a positive regulator of glutamatergic synapse development but does not influence GABAergic synapse development in the hippocampus.

**Figure 3:**
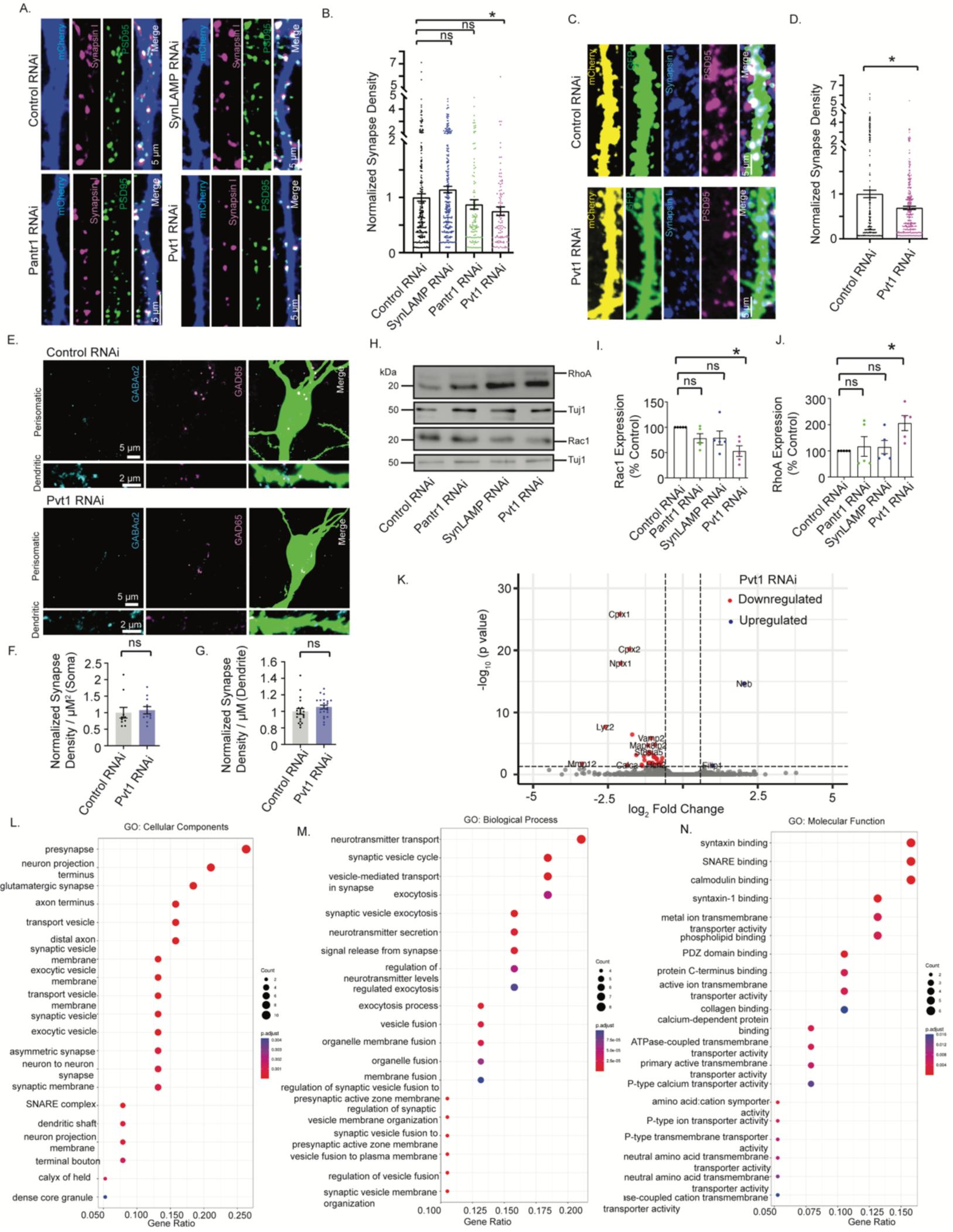
Pvt1 acts as a specific regulator of excitatory synapse development by influencing synaptogenic transcripts. **(A)** Images showing immunostaining of Synapsin I (magenta) and PSD95 (green) in the dendrites expressing mCherry (blue). Excitatory synapses are represented by apposed Synapsin I and PSD95 puncta on dendrites (white puncta). **(B)** Quantification of excitatory synapse density. n= 15–26 neurons from 3 independent experiments, *p<0.03, ns = not significant. One-way ANOVA and Fisher’s LSD. Data shown as mean ± SEM. (**C**) Excitatory synapse density analysis from *in utero* electroporated CA1 neurons expressing shRNA as indicated. Image showing immunostaining of Synapsin I (blue), PSD95 (magenta) on dendrites expressing mCherry (yellow) and EGFP (green). Overlapping PSD95 and Synapsin I puncta on dendrite represents glutamatergic synapse (white). **(D)** Quantification of excitatory synapse density. n= 36 – 39 dendrites from 10 - 12 neurons that are imaged from 3 independent experiments. *p<0.001. Unpaired t-test with Welch’s correction. Data shown as mean ± SEM. (**E**) Inhibitory synapse density analysis from *in utero* electroporated CA1 neurons expressing shRNA as indicated. Photomicrograph showing immunostaining of GABA-α1 (blue), GAD-65 (magenta) on soma and dendrites of CA1 neurons expressing EGFP (green). Overlapping GABA-α1 and GAD-65 puncta on soma and dendrites represent inhibitory synapses. **(F-G)** quantification of perisomatic and dendritic inhibitory synapse density. For perisomatic synapse density, n= 10-11 neurons; for dendritic synapse density, n= 20-23 dendrites from 3 independent experiments. ns= not significant. Unpaired t-test with Welch’s correction. Data shown as mean ± SEM. **(H)** Immunoblots of Rac1, RhoA and Tuj1 from cultured hippocampal neurons expressing shRNA as indicated. **(I-J)** Quantification of RhoA and Rac1 expression. n=5 independent experiments, *p< 0.02. ns = not significant, Unpaired t-test with Welch’s correction. Data shown as mean ± SEM. **(K)** Volcano plot showing up- or down–regulated coding transcripts after RNA-Seq analysis from Pvt1 RNAi neurons. n=3 independent experiments **(L-N)** GO analysis showing significant (FDR, p<0.05) enrichment of GO-terms for the function of genes encoding down-regulated mRNAs following Pvt1 RNAi. Also see Figure S4.

### Pvt1 influences the expression of cytoskeleton effectors and synaptic proteins

Regulators of the actin cytoskeleton determine the morphology of dendritic spines (Hotulainen and Hoogenraad, 2010; Kennedy et al., 2005). The Rho family of GTPases, such as RhoA, Rac1 and Cdc42, are key regulators of the actin cytoskeleton that profoundly influence spine morphology. Rac1 and Cdc42 promote spine growth, whereas RhoA enhances spine loss (Tolias et al., 2011). Prompted by these observations, we measured the expression of Rac1 and RhoA following the knockdown of Pantr1, SynLAMP and Pvt1. Hippocampal neurons (DIV 3-4) in culture were transduced with lentivirus expressing indicated shRNAs and Rac1 as well as RhoA protein levels were analyzed from these neurons (DIV 21). We observed that Pvt1 RNAi led to the reduction of Rac1 (47.00 ± 10.44 % decrease, p < 0.01), whereas the level of RhoA was enhanced (106.1 ± 28.89 % increase, p < 0.02) (Figure 3H - 3J). However, Pantr1 and SynLAMP RNAi had no effect (Figure 3H-3J).

To determine the Pvt1-driven molecular framework influencing synapse development, we next performed a transcriptomics analysis using total RNA from Pvt1 RNAi neurons. Cultured neurons (DIV 3-4) were transduced with lentivirus expressing shRNA against Pvt1 followed by RNA-seq analysis from these neurons (DIV 21) (Figure 3K). We found that Pvt1 RNAi predominantly reduced the expression of transcripts encoding post-synaptic proteins of glutamatergic synapses that include regulators of AMPA receptor activity, such as Nptx1; and presynaptic proteins involved in neurotransmitter transport such as VAMP2, Cplx1/2, Sk32a, Sv2c and Syt2. We also detected the upregulation of the actin binding protein Filip1, that negatively regulates dendritic spine morphology (Figure 3K). GO analysis of down-regulated transcripts (Table S2) revealed enrichment of synaptic GO annotations (FDR<0.05) (Figure 3L-3N). We also observed a significant enrichment in biological processes and cellular components associated with postsynaptic functions. The biological processes included, among others, neurotransmitter transport (p adj. 2.58944E-07), synaptic vesicle cycle (p adj. 3.83126E-06), vesicle-mediated transport to the synapse (p adj. 4.12905E-06), detection of stimulus (p adj. 0.000960904), protein localization to cell periphery (p adj. 0.002263423), and regulation of synaptic plasticity (p adj. 0.002299654). The cellular components include presynapse formation (p adj. 2.97269E-08), synaptic vesicle membrane proteins (p adj. 1.9457E-06), axon terminus (p adj. 3.20596E-06), glutamatergic synapse (p adj. 3.20596E-06), and synaptic vesicles (p adj. 8.61425E-05) (Figure 3L - 3N). There was also a significant enrichment of GO terms associated with molecular functions, such as syntaxin binding (FDR 8.279E-08), SNARE binding (FDR 6.62919E-07), and calmodulin binding ( FDR 8.58518E-06); all pointing towards Pvt1 having robust roles in the regulation of presynaptic and postsynaptic functions (Figure 3L-3N). This observation is consistent with our data showing that Pvt1 RNAi reduced the puncta density of both PSD95 (40.7978% decrease, p<0.03) and Synapsin I (37.6402% decrease, p<0.03) (Figures S4D - S4E).

### Regulation of AMPA receptor-mediated synaptic activity by lncRNAs

The recruitment of AMPA receptors (AMPARs) to the developing synapse drives the maturation of glutamatergic synapses. As synaptic transmission within the mature neural circuitry is predominantly governed by AMPAR activity (Hall and Ghosh, 2008; Wu et al., 1996), we used whole-cell voltage-clamp recordings to measure the amplitude and frequency of AMPAR–mediated miniature excitatory postsynaptic currents (mEPSCs) from cultured hippocampal neurons following the knockdown of each lncRNA. Neurons (DIV 6-7) were transduced with two types of lentivirus, each expressing a distinct shRNA (as indicated), and whole-cell voltage-clamp recording was performed from these neurons (DIV 21-25). We observed a significant reduction in mEPSC amplitude following the knockdown of SynLAMP (4.36 ± 0.37 pA decrease for shRNA#1, p<0.0001; 5.76 ± 0.30 pA decrease for shRNA# 2, p<0.0001), Pantr1 (4.33 ± 0.37 pA decrease for shRNA#1, p < 0.0001; 5.53 ± 0.32 decrease for shRNA#2, p<0.0001) and Pvt1 (5.35 ± 0.40 pA decrease for shRNA#1, p<0.0001; 5.67 ± 0.32 pA decrease for shRNA#2, p<0.0001) (Figures 4A - 4B, S5A - S5B). However, we did not observe any change in mEPSC frequency following SynLAMP or Pantr1 RNAi. (Figures 4A and 4C, S5A and S5C). A small reduction in mEPSC frequency was detected upon Pvt1 knockdown (1.67± 0.61 Hz decrease, p<0.007 for shRNA#1, 1.08 ± 0.39 Hz decrease, p<0.01 for shRNA#2) (Figures 4C and S5C). To counter confounding observations arising from the off-target effects of shRNAs, two independent shRNA–constructs against each lncRNA were used for the loss of function experiments.

**Figure 4:**
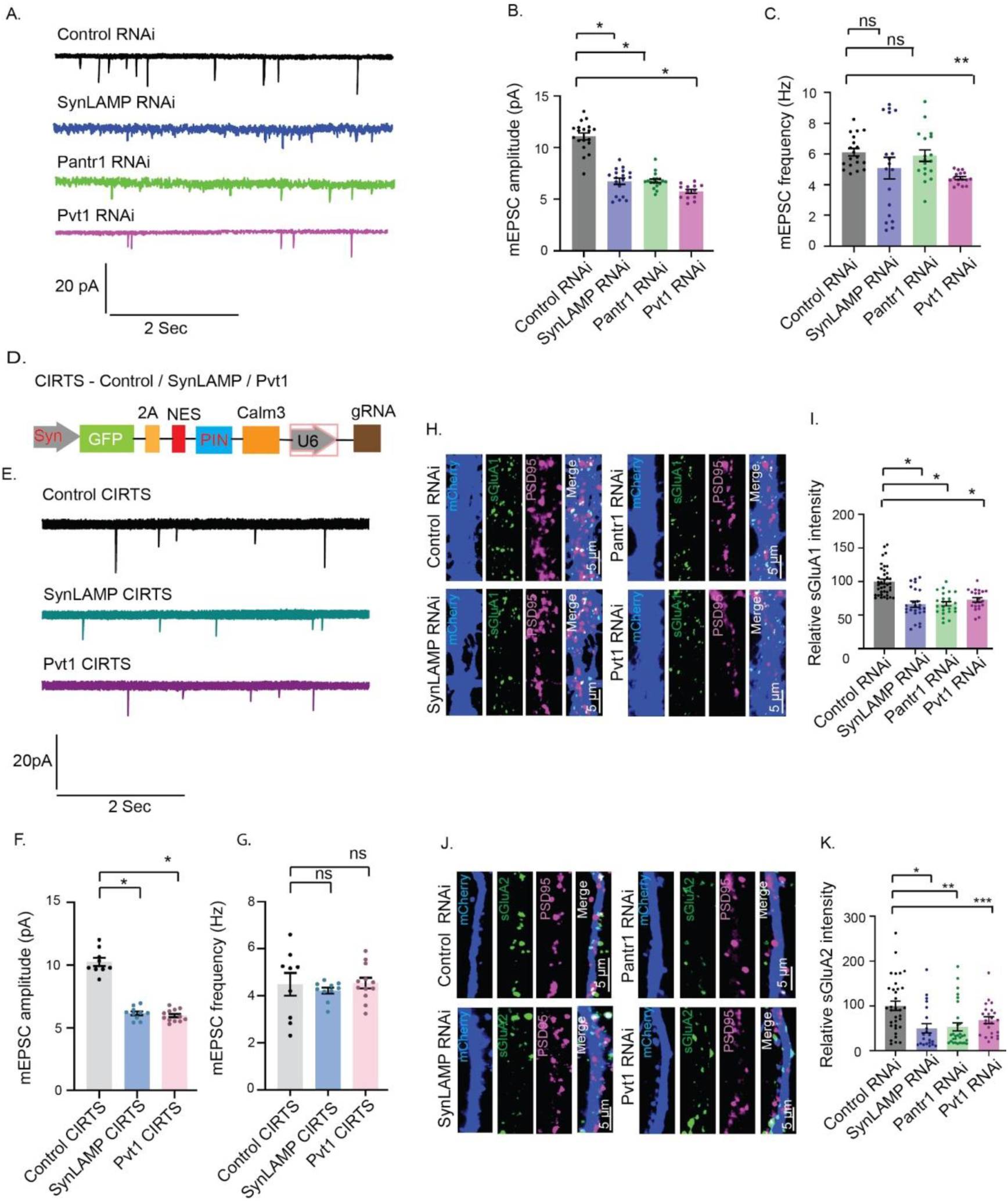
Loss of SynLAMP, Pantr1 or Pvt1 function affects synaptic activity *via* sAMPAR abundance. **(A)** mEPSC traces from whole-cell patch clamp recordings from hippocampal neurons with RNAi against indicated lncRNAs. **(B)** Mean mEPSC amplitude. **(C)** Mean mEPSC frequency. n = 16 – 19 neurons from 5 - 6 independent experiments. *p <0.0001, **p<0.007. ns = not significant. One-way ANOVA and Fisher’s LSD. Data shown as mean ± SEM. (**D**) Schematic representation of CIRTS construct. (**E**) mEPSC traces from whole-cell patch clamp recordings from hippocampal neurons with CIRTS-mediated knockdown of indicated lncRNAs. **(F)** Mean mEPSC amplitude. **(G)** Mean mEPSC frequency. n= 9 - 12 neurons from 3 independent experiments. One-way ANOVA and Fisher’s LSD. Data shown as mean ± SEM. **(H)** Photomicrograph showing sGluA1 abundance in dendrites (green puncta) following the knockdown of indicated shRNAs. Synapses marked by PSD95 (magenta puncta). sGluA1 abundance in synapses shown in merged (sGluA1/PSD95/mCherry) images. **(I)** Quantification of normalized intensity of synaptic sGluA1. n = 21 – 38 neurons from 3 independent experiments, *p< 0.0001. ns = not significant **(J)** Photomicrograph showing sGluA2 abundance in dendrites (green puncta) following the knockdown of indicated shRNAs. Synapses marked by PSD95 (magenta puncta). sGluA2 abundance in synapses shown in merged (sGluA2/PSD95/mCherry) images. **(K)** Quantification of normalized intensity of synaptic sGluA2. n = 21 – 34 neurons from 3 independent experiments, *p< 0.0005, **p<0.0004, ***p<0.03. ns = not significant. One-way ANOVA and Fisher’s LSD. Data shown as mean ± SEM. Also see Figure S5.

Further, to identify the synapse-specific roles of Pvt1 and SynLAMP, we selectively inhibited their function by a CRISPR-Cas-inspired-RNA-targeting system (CIRTS) and measured synaptic activity. The CIRTS system is dependent on a CALM3 intron that has a dendritic localization signal which spatially restricts the degradation of Pvt1 and SynLAMP in dendrites. Synaptic inhibition of either Pvt1 or SynLAMP significantly reduced mEPSC amplitude (Control CIRTS *vs* SynLAMP CIRTS: 4.086 ± 0.3087; p<0.0001; UC CIRTS *vs* SynLAMP CIRTS: 4.275 ± 0.2962, p<0.0001) without influencing the frequency (Figure 4D-4G). This observation indicates that synapse-localized lncRNAs can act in an autonomous manner to regulate synaptic activity.

mEPSC amplitude determines synaptic strength and is a direct readout of the abundance of AMPARs on the postsynaptic membrane (O’Brien et al., 1998; Srinivasan et al., 2021). The observed reduction in mEPSC amplitude therefore prompted us to measure AMPAR levels on the postsynaptic surface (sAMPARs) following knockdown of SynLAMP, Pantr1 and Pvt1. sAMPAR distribution was analyzed by measuring the surface expression of the GluA1 and GluA2 (sGluA1/A2) subunits of AMPARs. Cultured hippocampal neurons (DIV 6-7) were transduced with lentivirus expressing the indicated shRNA. Transduced neurons (DIV 21) were live labeled with N-terminus-specific antibodies against sGluA1/A2 and synapses were marked by PSD95. We found that sGluA1/A2 expression in excitatory synapses was reduced following the knockdown of SynLAMP (33.9 ± 5.1% decrease for sGluA1, p<0.0001; 50.11 ± 14.05% decrease for sGluA2, p<0.0005), Pantr1 (33.14± 5.17% decrease for sGluA1, p<0.0001; 46.87±12.79% decrease for sGluA2, p<0.0004) and Pvt1 (27.13 ± 5.39% decrease for sGluA1, p<0.0001; 31.97 ± 13.66 % decrease for sGluA2, p<0.03) (Figure 4H - 4K and S5D - S5E). Collectively, these findings indicate that SynLAMP and Pantr1 RNAi specifically affect the activity of mature synapses. The observed reduction in both frequency and amplitude following Pvt1 RNAi may be attributed to the formation of immature synapses (Figure 3–4).

### Contextual fear conditioning influences the synaptic distribution of lncRNAs

The localization of lncRNAs at the synapse prompted us to investigate the impact of synaptic activity on the dendritic distribution of lncRNAs. To achieve this, we used contextual fear conditioning (CFC), a hippocampus-based memory paradigm (Fanselow, 2000; Phillips and LeDoux, 1992) that relies on single-trial associative learning to assess the activity-induced distribution of lncRNAs in the dorsal hippocampus. The local abundance of lncRNAs was analyzed from synaptoneurosomes (SN) prepared from the dorsal hippocampus after measuring the freezing response 3h post training as the activity-induced translation of plasticity related proteins occur within this timeframe (Bourtchouladze et al., 1998; O’Donnell and Sejnowski, 2014; Sutton and Schuman, 2006). qRT-PCR analysis revealed that the abundance of SynLAMP (1.83 ± 0.3 fold, p<0.0009), Pantr1 (0.49 ± 0.09 fold, p<0.002) and Pvt1 (0.88 ± 0.14 fold, p<0.001) in the synaptic compartment was enhanced following CFC. Snhg8 was depleted (0.77 ± 0.13 fold decrease, p<0.001) from the synapse; whereas the distribution of 2900052N01Rik, Snhg12, Rmst1, Snhg6, Epb41l4aos and Zfas1 was not affected (Figure 5A-B). In order to determine whether the differential abundance of lncRNA was due to the altered expression of lncRNAs or due to the activity-dependent altered transport between the synapse and the cell body, we assessed lncRNA expression in the whole hippocampus and in the dorsal hippocampus 3 hours post-CFC.

**Figure 5:**
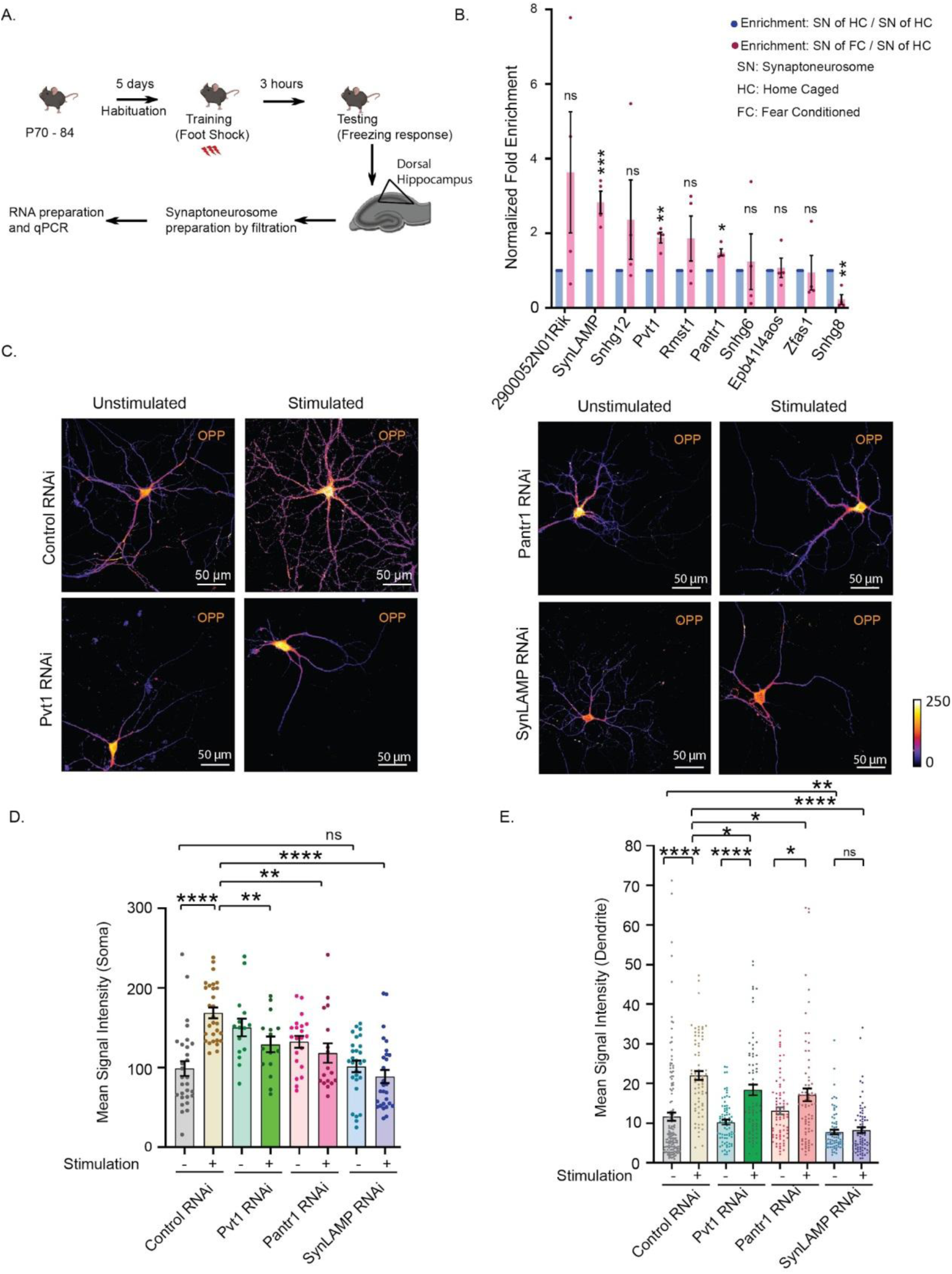
Activity-driven redistribution of synaptic lncRNAs and their role in regulating translation. (A) Schematic representation of contextual fear conditioning in mice and subsequent analysis of lncRNA expression from the SN fraction. (**B**) Synaptic abundance of indicated lncRNAs 3 hours post fear-conditioning. Histogram represents the relative abundance of lncRNAs. n = 4 independent experiments. *p< 0.002, **p<0.001, ***p<0.0009, ns = not significant. Multiple t-test determined by Holm-Sidak method. Data shown as mean ± SEM. (**C**) Images representing extent of activity-dependent translation in cultured neurons, transduced with shRNA (as indicated). Pixel intensity represented as a heat map.Calibration bar and scale bar as indicated. (**D-E**) Quantification of activity-induced translation in (**D**) soma and in (**E**) dendrites. n=15 neurons from 3 independent experiments. *p<0.05, **p<0.003, ****p<0.0001. ns= not significant. Two-tailed t-test with Welch’s correction. Data shown as mean ± SEM. Also see Figure S6 and S7.

There was a reduction in SynLAMP (0.29 ± 0.08 fold, p<0.01) and Pantr1 (0.37± 0.11, p<0.01) in the total hippocampus (Figure S6A). However, lncRNA expression levels remained unaltered in the dorsal hippocampus (Figure S6B). This dichotomy in lncRNA expression pattern suggests that CFC affects lncRNA stability within the hippocampus, but is restricted to areas outside the dorsal hippocampus. It would be interesting to study how lncRNAs in the ventral hippocampus and the dentate gyrus get affected due to CFC.

### Activity-dependent dendritic protein synthesis by lncRNAs

Enhanced abundance of SynLAMP, Pantr1 and Pvt1 in SN fraction following CFC prompted us to test their involvement in *de-novo* translation. Activity-dependent somatic and dendritic translation was scored by FUNCAT assay in glutamate-stimulated neurons that expressed shRNAs against SynLAMP/Pantr1/Pvt1 and dendrites were visualized by MAP2 immunostaining. Translation assays were performed in the presence of the transcription inhibitor (Actinomycin D, 10 μg/ml) to prevent the influence of nascent transcription. We observed that stimulation of Control RNAi neurons enhanced new translation both in the soma and the dendrites (soma: 69.87 ± 11.52 mean signal intensity (m.s.i) increase; p<0.0001; dendrites: 10.42 ± 1.52 m.s.i increase; p<0.0001; as compared to stimulated control RNAi neurons) while, SynLAMP RNAi led to reduced translation in both (soma: 79.75 ± 10.65 m.s.i decrease, p <0.0001; dendrites: 13.80 ± 1.32 m.s.i decrease, p<0.0001; as compared to stimulated control RNAi neurons) (Figures 5C-5E, S7A-S7B). The loss of Pvt1 or Pantr1 also reduced activity-regulated translation in both the soma and dendrites as compared to stimulated, control RNAi neurons (soma: 39.60 ± 11.90 m.s.i decrease in Pvt1 RNAi neurons, p<0.003; 50.40 ± 13.96 m.s.i decrease in Pantr1 RNAi neurons, p<0.003; dendrites: 3.630 ± 1.74 m.s.i decrease in Pvt1 RNAi neurons, p<0.05; 4.90 ± 1.95 m.s.i decrease in Pantr1 RNAi neurons, p<0.05); but the extent of reduction in protein synthesis following SynLAMP RNAi was more pronounced (Figures 5D-5E, S7A-S7B). Interestingly, basal translation within dendritic compartments was significantly hampered upon SynLAMP’s loss of function, while somatic translation remained unchanged (soma: 2.713 ± 11.76 m.s.i decrease w.r.t unstimulated control RNAi neurons, p<0.10; dendrites: 3.866 ± 1.187 m.s.i decrease as compared to unstimulated Control RNAi neurons, p<0.003). There was no significant change to the existing translational status of SynLAMP RNAi neurons upon stimulation (Figure 5D-5E).

### Synaptic activity influences the association of lncRNAs with RNA binding proteins

We wanted to identify how lncRNAs were being actively transported to the synapse following CFC. *De novo* translation at the synapse in the context of long-term memory formation relies substantially on RNA transport from the soma to dendritic spines. Transcripts in association with RNA binding proteins (RBPs) and translation regulators are packaged into “RNA granules” that are transported along the dendrites (Kiebler and Bassell, 2006). The activity-dependent release of RNA and associated translation factors from RNA granules promotes their transition from a translationally silent to an actively translating state (Krichevsky and Kosik, 2001; Shiina et al., 2005). Accordingly, we investigated the association between synapse-enriched lncRNAs and RBPs in response to CFC. We have analyzed the interaction between synapse-enriched lncRNAs and neuronal RBPs in a two-step process. First, we made an *in silico* interaction map based on the POSTAR3 (Zhao et al., 2021; Zhu et al., 2019) database that listed transcriptomics analysis from immunoprecipitated RBPs (Figures 6A and S8A).

**Figure 6:**
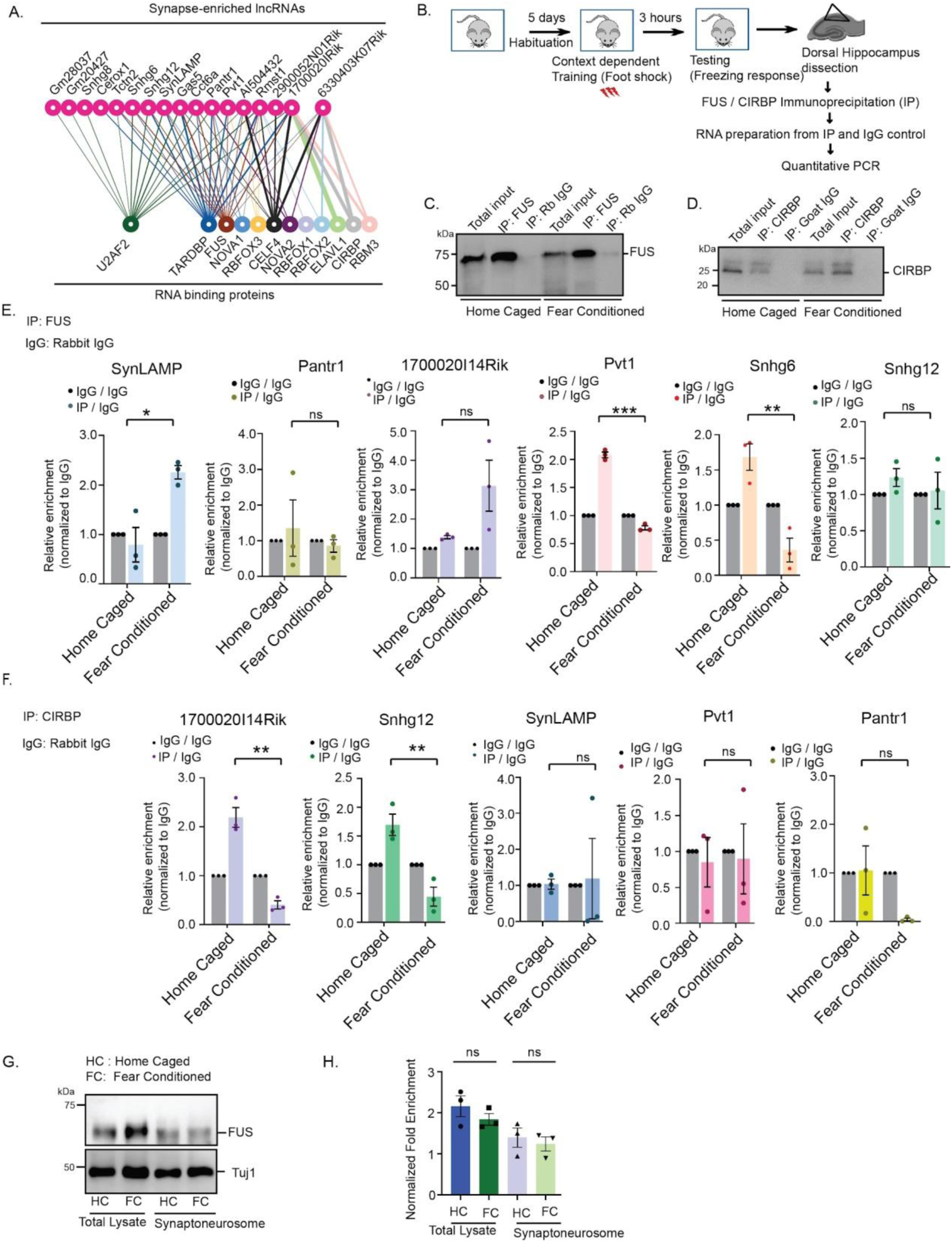
Activity-dependent interaction between synaptic lncRNAs and RBPs. (**A**) Schematic representation of synaptic lncRNA and RBP interactions based on PhyloP score reported in POSTAR3 database. (**B**) Paradigm used to identify activity-dependent association between RBPs and lncRNAs *in-vivo*. **(C-D)** Photomicrograph showing FUS **(C)** and CIRBP **(D)** immunoprecipitation from the dorsal hippocampus of adult mice upon 3 hours post contextual fear conditioning. (**E**) Association of indicated lncRNAs with FUS upon contextual fear conditioning. n=3 independent experiments. ***p<0.0001, **p< 0.007, *p< 0.04, ns= not significant. Unpaired t-test with Welch’s correction. Data shown as mean ± SEM. (**F**) Association of indicated lncRNAs with CIRBP upon contextual fear conditioning. n=3 independent experiments. **p<0.008, ns= not significant. Unpaired t-test with Welch’s correction. Data shown as mean ± SEM. Also see Figure S8. **(G)** Photomicrographs of FUS immunoblots from total tissue lysates and synaptoneurosome fractions from the dorsal hippocampus of adult mice following CFC. **(H)** Quantitation of **(G)**. ns= not significant. n=3 independent experiments. Data shown as mean ± SEM.Two-way ANOVA with FIsher’s LSD. Also See Figure S8

We assessed the number of binding motifs of specific RBPs present in a subset of synapse-enriched lncRNAs and evaluated their interaction based on the PhyloP score reported in POSTAR3 (Figure 6A). FUS (FUsed in Sarcoma) and CIRBP (Cold Inducible RNA Binding Protein) were immunoprecipitated from the dorsal hippocampus 3h post CFC (Figure 6B) and their interactions with lncRNAs were analysed (Figure 6C-6F). Fear conditioning led to a reduction in the association between FUS and Pvt1 (1.30±0.06 reduction in relative enrichment w.r.t IgG control, p<0.0002) or Snhg6 (1.32 ± 0.25 reduction in relative enrichment w.r.t IgG control, p<0.001), whereas, the association between FUS and SynLAMP was enhanced (1.46 ± 0.37 increase in relative enrichment w.r.t IgG control) (Figure 6E). This paradigm did not influence the interaction between FUS and Pantr1, Sngh12 and 1700020I14Rik. CFC reduced the association between CIRBP and 1700020I14Rik (1.79 ± 0.22 reduction in relative enrichment w.r.t IgG control, p<0.007) and Snhg12 (1.25±0.25 reduction in relative enrichment w.r.t IgG control, p<0.008). Although the PhyloP score of interaction between Snhg12 and CIRBP was not very high, it does have one binding site to CIRBP (Figure S8A). We did not detect any interaction between CIRBP and SynLAMP/Pantr1/Pvt1 as predicted by our *in silico* analysis (Figure 6F). A possibility of increased association between FUS and SynLAMP could be the simultaneous transport of FUS to the synapse. However, FUS levels remained comparable in the synaptoneurosomes of home-caged (control) *vs* fear-conditioned mice (Figure 6G - 6H). Together, these findings suggest two things; the existence of activity-dependent, reversible interactions between specific lncRNAs and RBP; and secondly, the possibility that neuronal activity allows SynLAMP to target synaptically localized FUS.

### Synaptic activity drives SynLAMP-dependent translation of the FUS target CamK2a

We found that CFC enhances the abundance of SynLAMP at the synapse without increasing FUS abundance, and there is enhanced association between FUS and SynLAMP upon neuronal activity. A plausible causality for supporting SynLAMP’s role as a molecular decoy is that neuronal activity allows SynLAMP to actively sequester synaptically-localized FUS. Since FUS is a translational repressor, sequestration of FUS should allow FUS-target mRNAs to be released from translation inhibition and be synthesized. To confirm this hypothesis, we performed proximity labelling assay (PLA) along with puromycin incorporation (Puro-PLA) to identify the nascent synthesis of CamK2a following the CIRTS-mediated knockdown of SynLAMP at the synapse (Dieck et al., 2012; Tom Dieck et al., 2015). While there was an expected increase in the Puro-PLA signal for CamK2a synthesis (fluorescence intensity) in the control group (100.8±30.56% increase, p<0.002); the observed increase was significantly abrogated upon SynLAMP knockdown at the synapse (67.03± 31.92% decrease, p<0.05) (Figure 7A-7B, S8B-8C) Thus, SynLAMP RNAi results in the inhibition of CamK2a translation inspite of glutamate stimulation and unaltered FUS levels.

**Figure 7:**
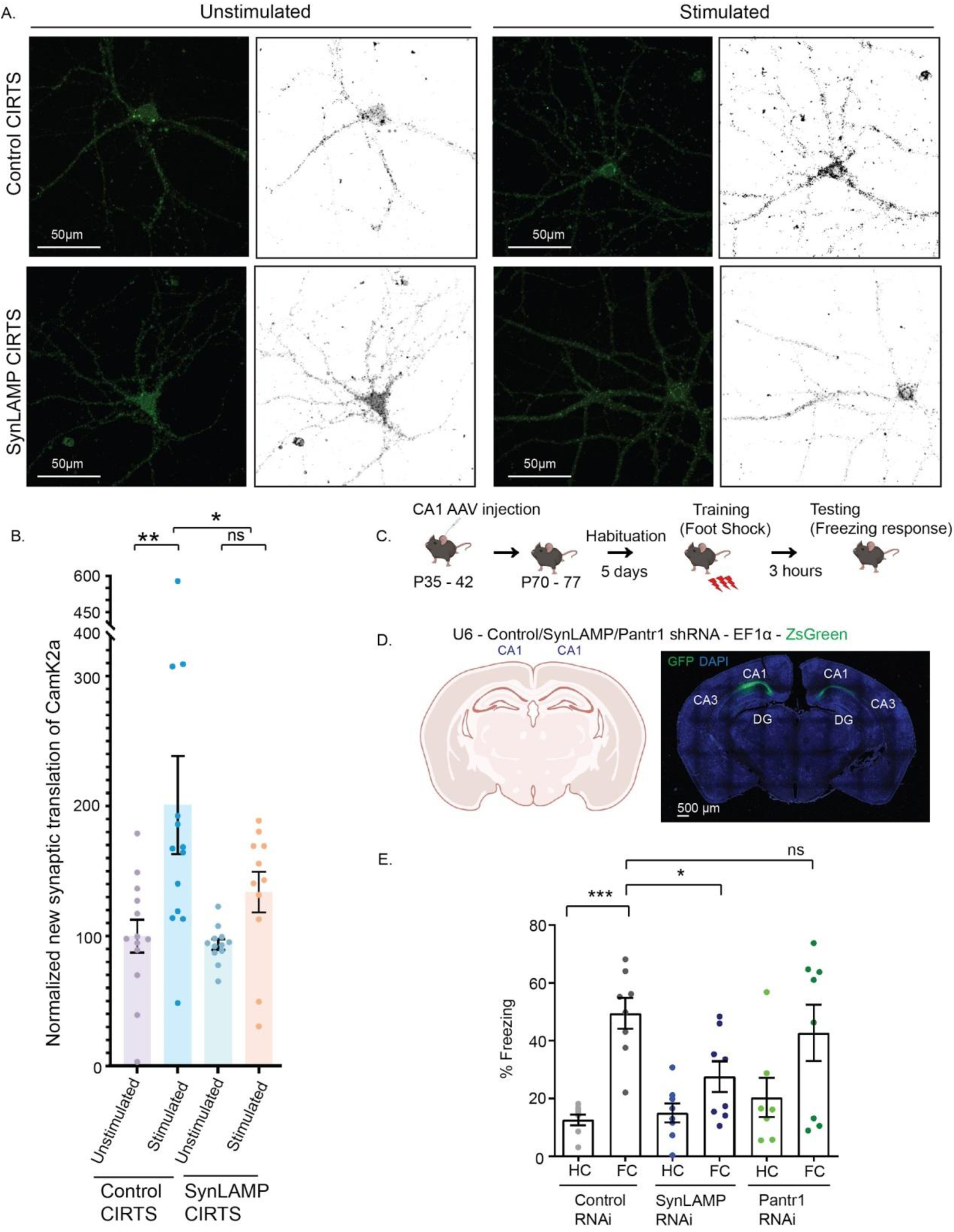
SynLAMP RNAi leads to deficits in contextual fear memory. (**A**) Photomicrographs of Puro-PLA assay in cultured hippocampal neurons with CIRTS-dependent knockdown of SynLAMP. Grey-scale images showing newly synthesised CamK2a puncta in basal condition or glutamate stimulation. Scale as indicated. **(B)** Quantitation of newly synthesized CamK2a puncta. n=11-13 from 3 independent experiments. **p<0.002, * p<0.05, ns= not significant. Data shown as mean ± SEM. One-way ANOVA with Fisher’s LSD. Also See Figure S8. **(C)** Contextual fear memory test paradigm following knockdown of SynLAMP/Pantr1 in CA1 neurons. **(D)** Schematic representation and photomicrograph of injected AAV constructs expressing shRNAs as indicated along with ZsGreen in CA1 neurons. Co-expression of ZsGreen in CA1 indicating effective manipulation of SynLAMP or Pantr1. **(E)** Contextual freezing response during test at 3 hours after conditioning following SynLAMP/Pantr1 RNAi. n=8, ***p<0.0001, *p<0.02. ns= not significant.Two way ANOVA with Fisher’s LSD. Data shown as mean ± SEM.

### SynLAMP (2410006H16Rik) is involved in contextual fear memory formation

The enrichment of SynLAMP and Pantr1 in CA1 dendrites following CFC and the significant reduction of glutamate-induced dendritic translation in SynLAMP and Pantr1 RNAi neurons prompted us to investigate their role in the regulation of fear memory. Adeno-associated virus (AAV) expressing ZsGreen with indicated shRNAs were injected into the CA1 of 7-8 week old mice. CFC was performed at 28-30 days later (Figure 7C-7E). We analyzed fear memory formation by measuring the freezing response 3h after training; this time window was selected on the basis of previous studies which report that application of translation inhibitors between 1h and 3h after training disrupts memory formation (Bourtchouladze et al., 1998; Sutton and Schuman, 2006). Although an expected increase in freezing between home-caged and fear-conditioned Control mice (36.88 ± 8.254 increase in % freezing, p< 0.0001) was observed, SynLAMP RNAi led to a reduction in freezing compared to the Control RNAi mice (difference: 21.53 ± 7.59 % freezing, p<0.02). In contrast, Pantr1 RNAi had no effect (Figure 7E). These data indicate that SynLAMP RNAi is specifically implicated in contextual fear memory.

## Discussion

### Localization of lncRNAs at the hippocampal synapse

Almost all lncRNAs characterized in the brain thus far localize to the nucleus. For example, the expression of Malat1, a lncRNA that is highly expressed in the brain, is thought to be restricted to the nucleus (Statello et al., 2021; Bernard et al., 2010). *neuroLNC* has been shown to influence neuritogenesis and neuronal migration *via* its interaction with the RBP TDP-43 (Keihani et al., 2019). A long nucleolus-specific lncRNA, (LoNA) affects translation by inhibiting the ribosomal RNA transcription and ribosome biosynthesis, and is implicated in memory (Li et al., 2018), and a newly discovered nuclear lncRNA, ADRAM, epigenetically regulates NR4A2 and the formation of fear extinction memory (Wei et al., 2022).

In contrast, very few cytoplasmic lncRNAs have been characterized. ADEPTR (activity-dependent transported lncRNA) has been shown to suppress a cAMP-induced increase in spine density and spontaneous synaptic activity (Grinman et al., 2021), but little is known about the identity and function of synapse-enriched lncRNAs. The results of this study fill this void, and lay the foundation to investigate the topological diversity of lncRNAs in regulating synaptic translation and diverse cognitive functions at the resolution of the synapse.

Towards this aim, we identified 1031 lncRNAs in synaptoneurosomes derived from the hippocampus. Among these transcripts, 94 lncRNAs were significantly enriched in the synaptic compartment (Figure 1). We validated the relevance of our transcriptomic analysis by comparing the coding transcripts found in our dataset with that of a previous study (Cajigas et al., 2012). Detection of well-known synapse-enriched coding transcripts in our dataset further emphasizes the accuracy of our transcriptomic analysis (Figure S2A). The number of lncRNAs detected in the fraction was comparatively lower than that of mRNAs previously identified in the CA1 neuropil (Cajigas et al., 2012). This could be attributed to the fact that the expression levels of lncRNAs are typically lower than those of mRNAs (Mukherjee et al., 2016). LncRNAs are commonly hypothesised to function as snoRNA-reserves as they harbor snoRNA sequences within their transcripts (Pelczar and Filipowicz, 1998). SnoRNAs are typically localized within the nucleolus and can be structurally categorised into C/D box snoRNAs and H/ACA box snoRNAs, named SNORD and SNORA respectively (Dong et al., 2009; Mourtada-Maarabouni et al., 2008). However, recent evidences show that certain lncRNAs exhibit cytoplasmic behaviour that are functionally distinct from their roles as mere snoRNA-reservoirs, and they have significant presence in the cytoplasm; such as Zfas1 (Askarian-Amiri et al., 2011).To visualize whether synaptically-enriched lncRNAs in our list also possess snoRNA signatures, we have analyzed the snoRNA distribution in those transcripts. We observed that none of these synaptically enriched lncRNAs display signatures of embedded snoRNAs (Figure 1F). We further analysed the transcript distribution for a subset of synapse-enriched lncRNAs (Figure S2B). Isoforms of select lncRNAs are localized in the synapse; indicating that distinct isoforms of lncRNAs occupy different subcellular niches. Consistent with the existence of diverse classes of lncRNAs (Mattick, 2023), our annotated lncRNAs included intergenic, antisense and processed transcripts.

It would be worthwhile to explore in future studies, what are the unique signatures of the neuronally enriched novel lncRNAs and how they contribute to the regulation of neuronal functions.

### Pvt1 regulates cytoskeletal effectors of neuronal development

The roles of cytoplasmic lncRNAs in regulating dendritic arborization and spine morphology *in vivo* have not been investigated. Pvt1 knockdown led to a significant reduction in dendrite complexity and spine density, together with altered spine morphology. Furthermore, Pvt1 RNAi reduced stubby and mushroom spines *in vitro* and *in vivo*, whereas knockdown of SynLAMP and Pantr1 had no effect on either dendritic complexity or spine morphology (Figure 2). These findings emphasize the involvement of specific dendritic lncRNAs in spine development within the developing mouse brain.

To investigate the signaling mechanisms underlying Pvt1-driven dendritic growth and spine morphology, we explored the role of Rho-related GTPases. RhoA and Rac1 are expressed in hippocampal and cortical neurons during the window of dendrite growth and spine development (Nakayama and Luo, 2000; Schubert and Dotti, 2007; Threadgill et al., 1997). As there is reduction of dendritic growth (Threadgill et al., 1997) and spine density (Nakayama and Luo, 2000) in dominant negative Rac-1 phenotypes, the reduction in spine density and spine shrinkage observed in Pvt1 RNAi neurons (Figure 2) could be mediated by Rac-1 (Figure 3). The active form of RhoA reduces dendritic branching (Nakayama and Luo, 2000) and spine number (Schubert and Dotti, 2007. We observed the enhancement of RhoA expression following Pvt1 RNAi (Figure 2). Reduced dendrite growth and altered spine morphology with a concomitant increase in RhoA expression in Pvt1 RNAi neurons indicate that Pvt1 negatively regulates active forms of RhoA.

Transcriptomics analysis following Pvt1 knockdown did not show any change in RhoA and Rac1 transcripts (Figure 3); suggesting an indirect involvement of Pvt1 in the regulation of active RhoA and Rac1. Another dendritic lncRNA, ADEPTR, has been shown to regulate spine morphology in a cyclic AMP-Protein Kinase A (cAMP-PKA)-dependent manner (Grinman et al., 2021). Pvt1-mediated regulation of dendrite growth and spine morphology *via* Rho GTPases indicate the existence of a distinct pathway other than cAMP-PKA signaling.

### Pvt1 is a specific regulator of glutamatergic synapse formation

Our results revealed that Pvt1 RNAi reduced excitatory, but not inhibitory, synapse development in CA1 neurons (Figure 3). Pvt1 is the first example of an lncRNA that is implicated specifically in excitatory synapse development. Prior to our study, Malat1 has been shown to affect synapse density in cultured hippocampal neurons; however, this observation is primarily based on measuring the immunoreactivity of Synapsin I (Bernard et al., 2010). This data could be an indicator of pre-synaptic differentiation as observed in developing cerebellar synapses (Castejón et al., 2004). In contrast, our data confirmed the necessity of Pvt1 in excitatory synapse formation as we analyzed synapse density by measuring the co-localization of both pre and post-synaptic proteins *in vitro* and *in vivo*.

How does Pvt1 regulate excitatory synapse development? RNASeq and GO analysis from cultured neurons identified a number of transcripts that were down-regulated due to Pvt1’s loss of function (Figure 3). Most of the Pvt1-regulated transcripts were involved in glutamatergic synapse formation, presynapse, neuron projection, neurotransmitter transport and regulation of vesicle-mediated transport to the synapse; among others (Figure 3), as evident from the GO enriched terms. It is important to note that a transcription inhibitor was not used in both studies; therefore our transcriptomics analysis captured the differential regulation of mRNAs contributed by changes in transcription or mRNA stability.

### Synapse-enriched Pvt1 regulates the synaptic activity in mature synapses

Our patch-clamp experiments indicate that Pvt1 has significant influence on both the pre- and post-synapse in excitatory neurons. Observed changes in synaptic activity could either be due to Pvt1’s loss of function specifically in the synaptic compartments, or the effect of a pan-neuronal knockdown of the lncRNA. To explore these possibilities, we achieved dendrite specific knockdown of Pvt1, using a synapse-targeted CRISPR-based CIRTS construct (Liau et al., 2023), leaving the remaining cytosolic Pvt1 unaltered. Measurement of mEPSCs from these neurons showed a significant reduction in mEPSC amplitude without affecting mEPSC frequency. However, a pan-neuronal knockdown of Pvt1 resulted in the reduction of both mEPSC amplitude and frequency (Figure 4). A comparative analysis of these patch-clamp studies not only defines the involvement of Pvt1 in the regulation of synaptic activity, but also emphasizes that synapse-localized Pvt1 has contributions to synaptic functions that are distinct from its global role.

Coupled with the observed reduction in Synapsin I and PSD95 density in Pvt1 RNAi neurons (Figure S4-S4E), and the reduction in mRNAs associated with glutamatergic synapse development and transmission;our data strongly implicates Pvt1 involvement in various stages of excitatory synapse development and function.

### Synapse-enriched lncRNAs modulate surface AMPAR distribution in the postsynaptic compartment

Loss of Pantr1/SynLAMP function in neurons resulted in the concomitant reduction of mEPSC amplitudes and surface AMPA receptor (sAMPARs) distribution in the post-synaptic membrane (Figure 4). The noticeable absence of developmental defects in hippocampal neurons after SynLAMP and Pantr1 RNAi (Figure 2–4) indicate that these two lncRNAs are specifically required for the tuning of mature synapses and their function is predominantly synapse-centric. Pvt1 RNAi also shows a decrease in sAMPARs (Figure 4H-4K); however, taking into consideration multiple lines of evidence from the synapse formation assays and GO enrichment analysis (Figure 2–3), it is possible that the observed decrease in sAMPARs is due to the lesser number of synapses formed. Further research is required to determine the mechanistic details of sAMPAR regulation by lncRNAs.

### Pvt1 is involved in both synapse development and plasticity

Pvt1 regulates dendritic complexity and synapse maturation in the early stages of neuronal development (Figure 2–3); it also undergoes activity-induced transport to regulate translation at mature synapses (Figure 4–6). These two observations can be reconciled if the temporal resolution of lncRNA function at different stages of neuronal development is considered. Spatiotemporal compartmentalization of Pvt1 function can be achieved through its diverse isoforms, the expression of which may take precedence at specific timepoints. We detected at least ten isoforms of Pvt1 in our bulk RNA-seq (Figure S2). However, shRNAs designed against Pvt1 were not isoform specific; thus all phenotypes resulting from Pvt1’s loss of function were observed. Diverse isoforms of lncRNAs may possess distinct temporal signatures in neurons (Mercer et al., 2012) and recruit different molecular partners to regulate markedly different signaling modules during developmental decisions or synaptic plasticity.

### Somatodendritic distribution of lncRNAs are invoked by neuronal activity

The localization of coding and non-coding transcripts in dendrites create specialized microdomains that determine the availability of plasticity-related proteins for synaptic plasticity and memory. Neuronal activity has been shown to influence the distribution of transcripts at the synapse (Ho et al., 2011; Sutton and Schuman, 2006). We therefore analyzed the subcellular abundance of lncRNAs in CA1 neurons following CFC as learning shapes the translation state of CA1 pyramidal neurons (Klann and Dever, 2004; Sutton and Schuman, 2006). CFC is a very strong learning paradigm with the ability to invoke activity-induced changes within a large number of neurons *in vivo*. CFC enhanced the synaptic distribution of SynLAMP, Pantr1 and Pvt1, but reduced the abundance of Snhg8. The synaptic localization of Snhg6, Snhg12, Zfas1, Rmst1, Epb4Iaos and 2900052NO1Rik were not affected (Figure 5). These data suggest an activity-invoked, state-dependent, and transcript-specific distribution of lncRNAs in the synapse. SynLAMP, Pvt1 and Pantr1 are involved in various aspects of synapse maturation and the regulation of synaptic output. It may be speculated that the selective localization of lncRNAs to the synapse in response to neuronal activity allows them to influence the synaptic proteome by regulating the translation or post-translation modifications of localized coding transcripts. As we observed the localisation of the lncRNAs 3h after CFC; there is a high probability that they are involved in regulating protein synthesis at or near synapses immediately following a learning event.

### Synaptically distributed lncRNAs regulate activity-dependent dendritic translation

Activity-dependent localization of transcripts in dendrites is essential to meet the demand for site-specific translation. Non-coding RNAs act as independent effectors that regulate translation *via* diverse mechanisms (Mercer et al., 2009). Our understanding of non-coding RNA mediated translation in dendrites is primarily limited to miRNAs (Banerjee et al., 2009; Sambandan et al., 2017; Schratt et al., 2006). We found that there was a reduction in dendritic translation upon the individual knockdown of SynLAMP/ Pantr1/Pvt1; with SynLAMP RNAi having the most pronounced effect (Figures 6 and S6). Our finding is the first evidence supporting the involvement of synaptic lncRNAs in the regulation of activity-dependent translation.

### Neuronal activity reverses the association between synaptic lncRNAs and RBPs

Transport of SynLAMP/Pantr1/Pvt1/Snhg8 in somato-dendritic compartments of the dorsal hippocampus within 3h post CFC (Figure 5) and their implications in dendritic translation prompted us to explore lncRNA-RBP interactions following CFC. The activity-dependent transport of regulatory RNAs is associated with changes to their metabolic state and requires the participation of select RBPs (Fernandez-Moya et al., 2014; Heraud-Farlow et al., 2013; Schieweck et al., 2020). Indeed, RBPs are key determinants of RNA transport, stability and translation (Bramham and Wells, 2007).

Here, we represent the RBP-lncRNA network as a multigraph plot; wherein line density is commensurate to the predicted lncRNA-RBP affinity (Figure 6). We tested the lncRNA association with two RBPs; FUS and CIRBP. FUS localizes to CA1 dendrites (Belly et al., 2005), forms RNA granules to regulate *de novo* translation (Yasuda et al., 2013), and affects mRNA stability (Udagawa et al., 2015). Increased mRNA affinity of FUS has been shown to inhibit translation in an mTOR-dependent manner (Sévigny et al., 2020). CIRBP also localizes in the dendrites of hippocampal neurons (Tong et al., 2013; Zhou et al., 2021); and has been shown to regulate RNA stability (Logan and Storey, 2020; Xia et al., 2012) and translation (De Leeuw et al., 2007) in non-neuronal cells. While there was no change to the overall distribution of FUS in the synaptic compartments upon CFC, we detected enhanced synaptic abundance of SynLAMP. The increased interaction between SynLAMP and FUS upon CFC suggests that SynLAMP functions as a sponge to sequester the translation repressor. Conversely, we observed the CFC-invoked dissociation of Pvt1 from FUS (Figure 6). The sponge activity of SynLAMP is specific as we have detected the reduction in association between FUS and the lncRNAs Pvt1 and Snhg6; whereas 1700020I14Rik was unaffected. It is important to note that, the synaptic abundance of SynLAMP is far greater than Pvt1 based on our transcriptomic data (Figure 1). Therefore, the possibility of sequestering FUS by SynLAMP could play an overtly dominant role in the regulation of FUS activity.

In contrast, CIRBP interacted with 1700020I14Rik and their association was reduced upon CFC (Figure 6). Of note, 1700020I14Rik was associated with both FUS and CIRBP but synaptic activity only affected its interaction with CIRBP (Figure 6). These data suggest an activity-dependent and transcript-specific interaction of lncRNAs with FUS/CIRBP. Correlating this observation with our *in silico* prediction whereby a single lncRNA can bind with multiple RBPs, we speculate that lncRNAs could function as a decoy for RBPs (Hafezqorani et al., 2019; Ray et al., 2013) and synergistically determine mRNA stability or translation through the formation of RNA granules.

### Synapse-specific loss of SynLAMP regulates FUS-dependent translation

Memory consolidation following a learning paradigm, such as CFC relies on the increased recruitment of coding/noncoding transcripts at the synapse and concomitant increase in dendritic translation following training (Bourtchouladze et al., 1998; Frey and Morris, 1997; Montarolo, 1986). To delineate synapse-specific roles of SynLAMP in regulating synaptic activity, we employed a CRISPR-CIRTS construct to perform SynLAMP knockdown in synapses (Figure 4), and measured neuronal output. Synaptic loss of function of SynLAMP recapitulates the reduction in mEPSC amplitude observed upon the pan-neuronal RNAi of SynLAMP, suggesting that SynLAMP’s functions are restricted to the post-synaptic compartment. This observation, along with the enhanced association of SynLAMP and FUS upon CFC, prompted us to explore possible molecular decoy properties of the lncRNA. Using Puro-PLA assay in SynLAMP-CIRTS neurons, we found that SynLAMP knockdown partially abrogates the stimulation-dependent translation of the FUS target CamK2a. CamK2a is known to be translated in synaptic compartments in an activity-dependent manner during memory consolidation (Miller et al., 2002) and FUS is its translation repressor localised to the synaptic compartment (Sahadevan et al., 2021). Collectively, our data provides a proof-of-principle evidence wherein activity-dependent localization of SynLAMP regulates the local synthesis of CamK2a by acting as a RNA-decoy to the RBP FUS.

### The synaptic lncRNA SynLAMP regulates contextual fear memory

Long-term memory formation is regulated by the activity-induced distribution of RNAs in dendrites (Swarnkar et al., 2021) and *de novo* translation from plasticity-related transcripts at the synapse (Ho et al., 2011). We observed that loss of SynLAMP in CA1 neurons disrupted contextual fear memory (Figure 7). Although Pantr1 is enriched in CA1 neurons (Goff et al., 2015) and positively regulates translation and mEPSC amplitudes in mature neurons; Pantr1 knockdown in CA1 neurons had no effect on contextual fear memory (Figure 7). We therefore speculate that Pantr1 could be involved in other forms of protein synthesis–dependent plasticity (Srinivasan et al., 2021) rather than Hebbian plasticity. As SynLAMP functions as a positive regulator of activity-dependent translation (Figure 5), and SynLAMP RNAi disrupts memory consolidation, SynLAMP–mediated dendritic protein synthesis could be a link to contextual fear memory (Figures 5 and 7).

In summary, we provide conclusive evidence for the localization and function of lncRNAs in the synaptic compartment, *in vivo*. It is evident that lncRNAs are critically involved in synaptic plasticity and experience-dependent regulatory activities linked to diverse cellular pathways.This study further opens up avenues to explore localized roles of lncRNAs in the regulation of synaptic translation. This discovery adds a new dimension to the RNA-centric view of developmental and cognitive functions of the brain, and to their dysfunction during neurodevelopmental disorders.

## Supplementary Information

**Figure S1:**
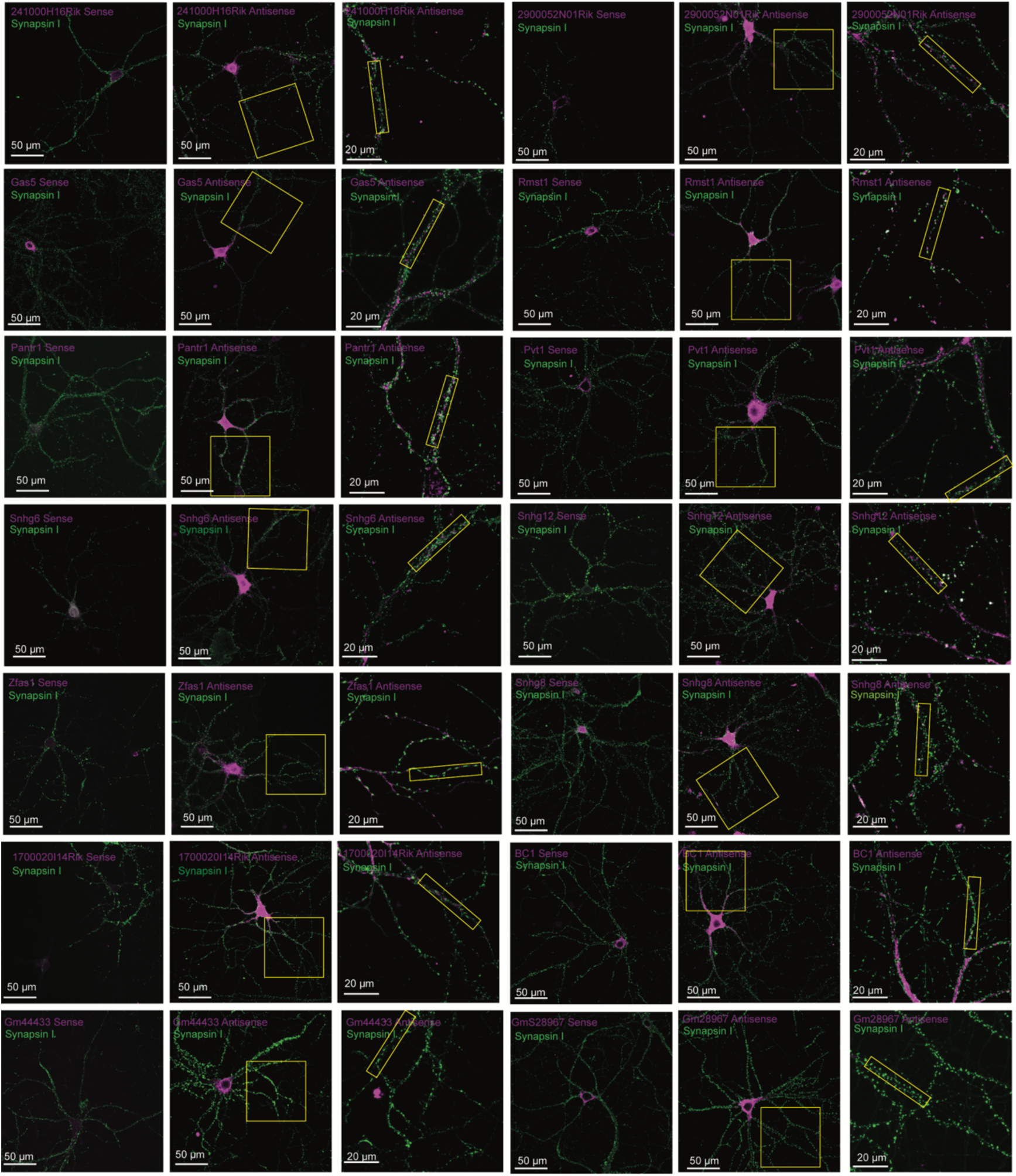
*In situ* hybridization of indicated synapse-enriched lncRNAs. Cultured hippocampal neurons (DIV21) were hybridized with indicated sense and antisense probes (magenta). Synapses are marked with Synapsin I (green puncta). High magnification images of dendrites that are shown in Figure 1 are marked in “Yellow” box. Scale as indicated. Also see Figure 1.

**Figure S2:**
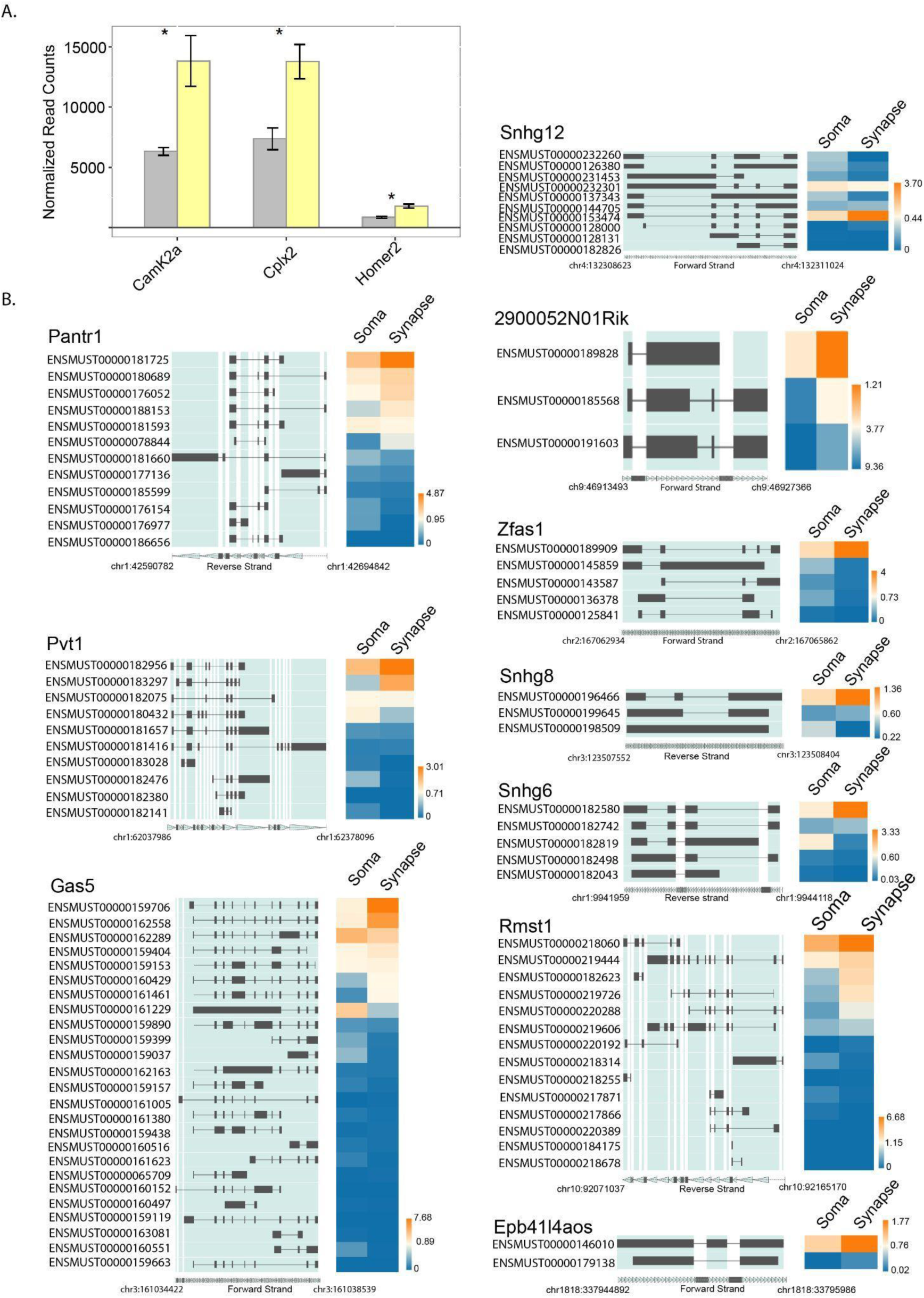
Spliced isoforms of lncRNAs that are enriched at the synapse. **(A)** Comparison of coding transcripts between our dataset with those previously published revealed similar enrichment patterns of coding transcripts between the soma ( grey ) and synapse (yellow). **(B)** Spliced isoforms of lncRNAs that are abundant at the synapse. Scale and chromosomal location as indicated.

**Figure S3:**
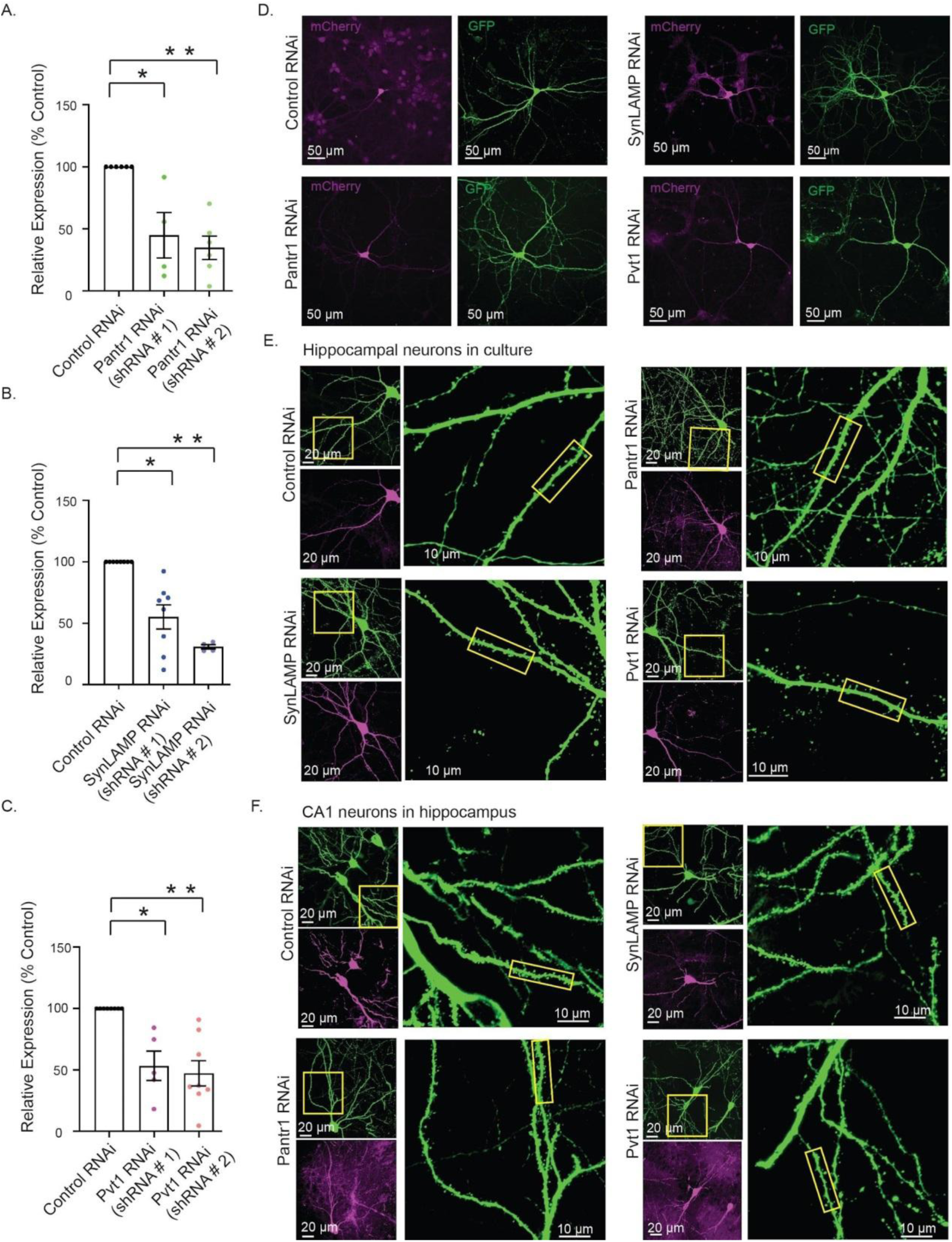
Spine development analysis following knockdown of Pantr1, SynLAMP and Pvt1. (**A-C**) Cultured hippocampal neurons (DIV 5 – 7) were transduced and total RNA was purified from these neurons (DIV 21) for qPCR analysis. Tuj1 expression was used for normalization of respective lncRNA expression. (A) Knockdown of Pantr1 by two distinct shRNAs, *p<0.005 for shRNA#1, **p< 0.0001 for shRNA#2 (B) Knockdown of SynLAMP by two distinct shRNAs, *p<0.0004 for shRNA#1, **p<0.0001 for shRNA#2. (C) Knockdown of Pvt1 by two distinct shRNAs.*p<0.0003 for shRNA#1, **p<0.0002 for shRNA2. 4-8 independent experiments. Unpaired two-tailed t-test. Data shown as mean ± SEM. (**D-E**) Cultured hippocampal neurons (DIV 3-4) were co-transfected with plasmids expressing indicated shRNAs along with mCherry and EGFP. (D) Cytosolic EGFP in neurons (DIV 21) was used to trace dendrites for Sholl analysis. (E) Cytosolic EGFP in neurons (DIV21) was used for dendritic spine analysis. High magnification images are indicated in “yellow” box. Scale as indicated. Also see Figure 2. (**F**) CA1 neurons were co-electroporated (E17 embryos) with plasmids expressing mCherry along with indicated shRNAs and EGFP. Spines were analyzed from CA1 neurons (P28). High magnification images are indicated in “yellow” box. Scale as indicated. Also see Figure 2.

**Figure S4:**
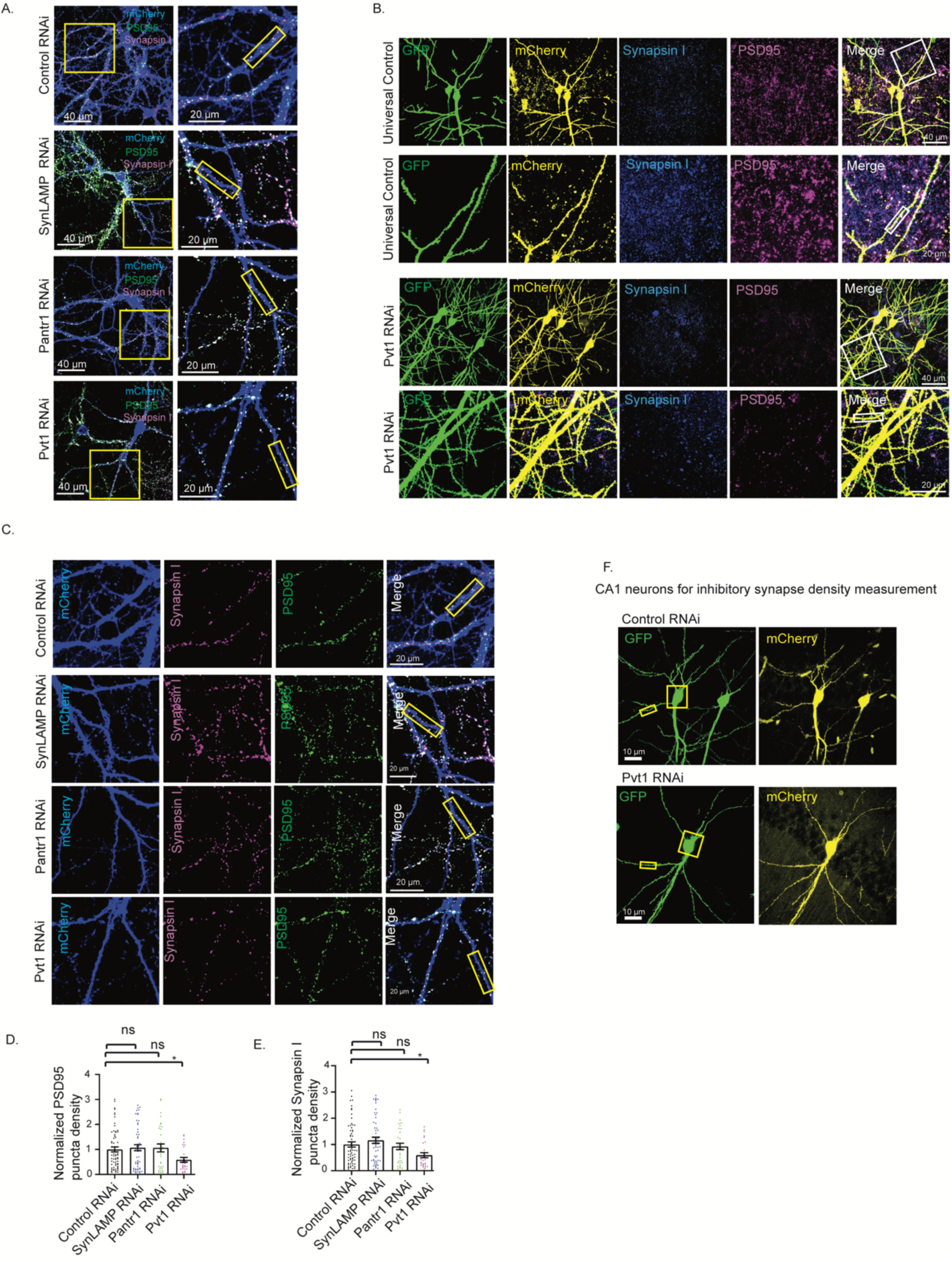
Effect of Pvt1 knockdown in synapse development. (**A**) Cultured hippocampal neurons (DIV 4-5) were transduced with lentivirus expressing control shRNA or shRNA against Pantr1/SynLAMP/Pvt1 (as indicated). Images from neurons (DIV21) showing PSD95 (green), Synapsin I (magenta) and dendrites (blue). Glutamatergic synapses (white) are represented by overlapping PSD95 and Synapsin I puncta on dendrite. Scale as indicated. High magnification images were captured from dendrites (yellow).(**B**) CA1 neurons (at E17) were *in utero* electroporated with plasmids expressing mCherry along with shRNA against Pvt1 or control shRNA and EGFP. Images of neurons (P28) showing PSD95 (magenta), Synapsin I (blue) and dendrite (mCherry and EGFP). Glutamatergic synapses (white) showing apposing PSD95 and Synapsin I puncta. High magnification images were captured from dendrites (yellow and green). Scale as indicated. White box indicates the ROI represented in Figure 3. (**C**) Representative image from cultured hippocampal neurons showing PSD95 and Synapsin I levels in neurons transduced with lentivirus expressing indicated shRNA. (**D**) Quantification of PSD95 puncta density from cultured neurons. n = 15 - 26 from 3 independent experiments, *p<0.03. ns = not significant. One-way ANOVA and Fisher’s LSD. Data shown as mean ± SEM.(**E**) Quantification of Synapsin I puncta density from cultured neurons. n = 15 - 26 from 3 independent experiments, *p<0.03. ns = not significant. One-way ANOVA and Fisher’s LSD. Data shown as mean ± SEM. (**F**) Image showing CA1 neurons expressing control shRNA and Pvt1 shRNA. Inhibitory synapse density onto these neurons were measured. Yellow boxes indicate regions of neurons represented in panels of Figure 3E-3F. Scale as indicated. Also see Figure 3.

**Figure S5:**
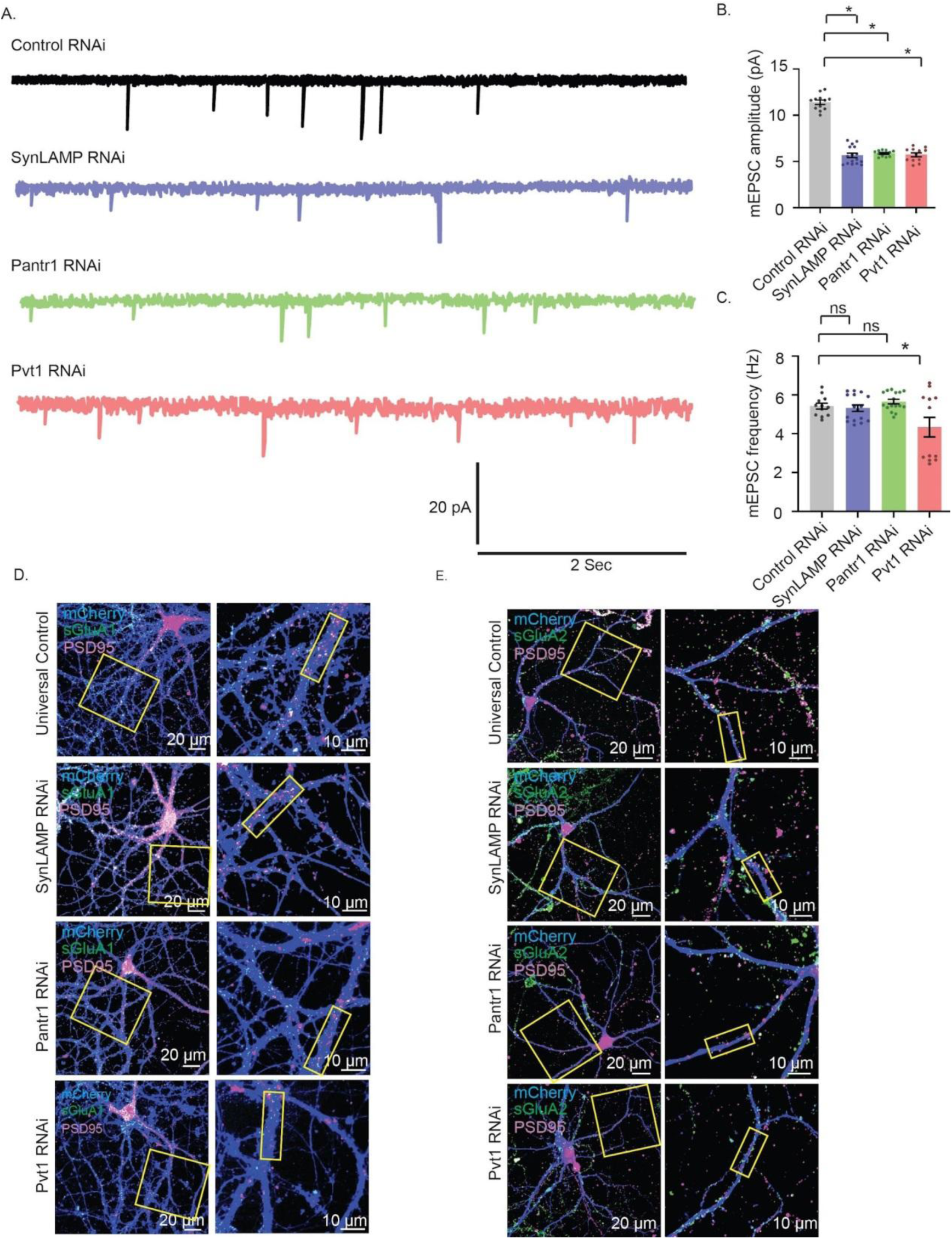
SynLAMP, Pantr1 and Pvt1 RNAi influence synaptic activity *via* sAMPAR abundance. (**A – C**) Cultured hippocampal neurons (DIV 10) were transduced with lentivirus expressing shRNAs#2 as indicated. Whole-cell patch clamp recordings were performed from transduced neurons (DIV 21 - 25). (A) mEPSC traces. (B) Mean mEPSC amplitude. (C) Mean mEPSC frequency. n = 12 – 16. *p <0.01. ns = not significant. One-way ANOVA and Fisher’s LSD. Data shown as mean ± SEM. See Figure 4 for shRNA#1 data point.(**D – E**) Cultured hippocampal neurons (DIV 10) were transduced with lentivirus expressing shRNAs # 1 as indicated. sAMPAR abundances were measured by immunolableing for GluA1 or GluA2 of live transduced neurons (DI V21 - 23). Following GluA1 or GluA2 labelling neurons were fixed and immunostained with PSD95 to mark synapses. Image showing (D) sGluA1 (green) or (E) sGluA2 (green) on mCherry expressing dendrites (blue) from cultured neurons immunostained with PSD95 (magenta). sGluA1 or sGluA2 abundance in synapses shown in merged (sGluA1 or sGluA2/PSD95/mCherry) images. Yellow box indicates the ROI represented in Figure 4. See also Figure 4.

**Figure S6:**
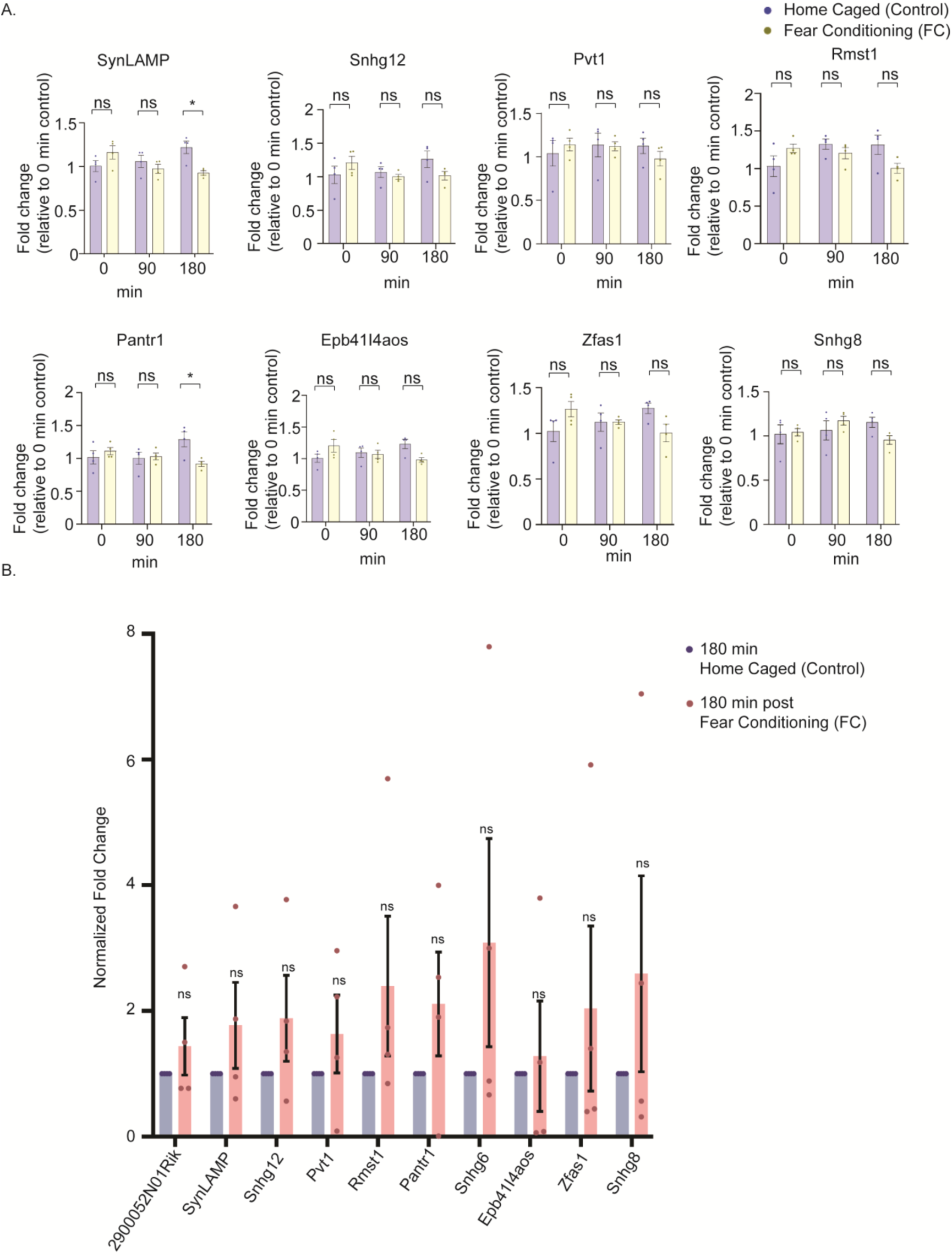
Expression of lncRNAs in hippocampus upon contextual fear conditioning. (**A**)Total RNA from the whole hippocampus was purified at 0 minutes, 90 minutes and 180 minutes following contextual fear conditioning. Indicated lncRNAs were analyzed by qRT-PCR. Bar plots represent relative fold change of indicated lncRNAs.*p<0.01, ns = not significant. n = 4, 2 Way ANOVA and Bonferroni multiple comparison test. Data shown as mean ± SEM.(**B**) The dorsal hippocampus (majorly CA1 region) was isolated from adult mice three hours (180 minutes) post contextual fear conditioning. Abundance of indicated lncRNAs was analyzed by qRT-PCR subsequent to total RNA isolation from the dorsal hippocampus. n = 4, ns = not significant. Unpaired two tailed t-test. Data shown as mean ± SEM. Also see Figure 5.

**Figure S7:**
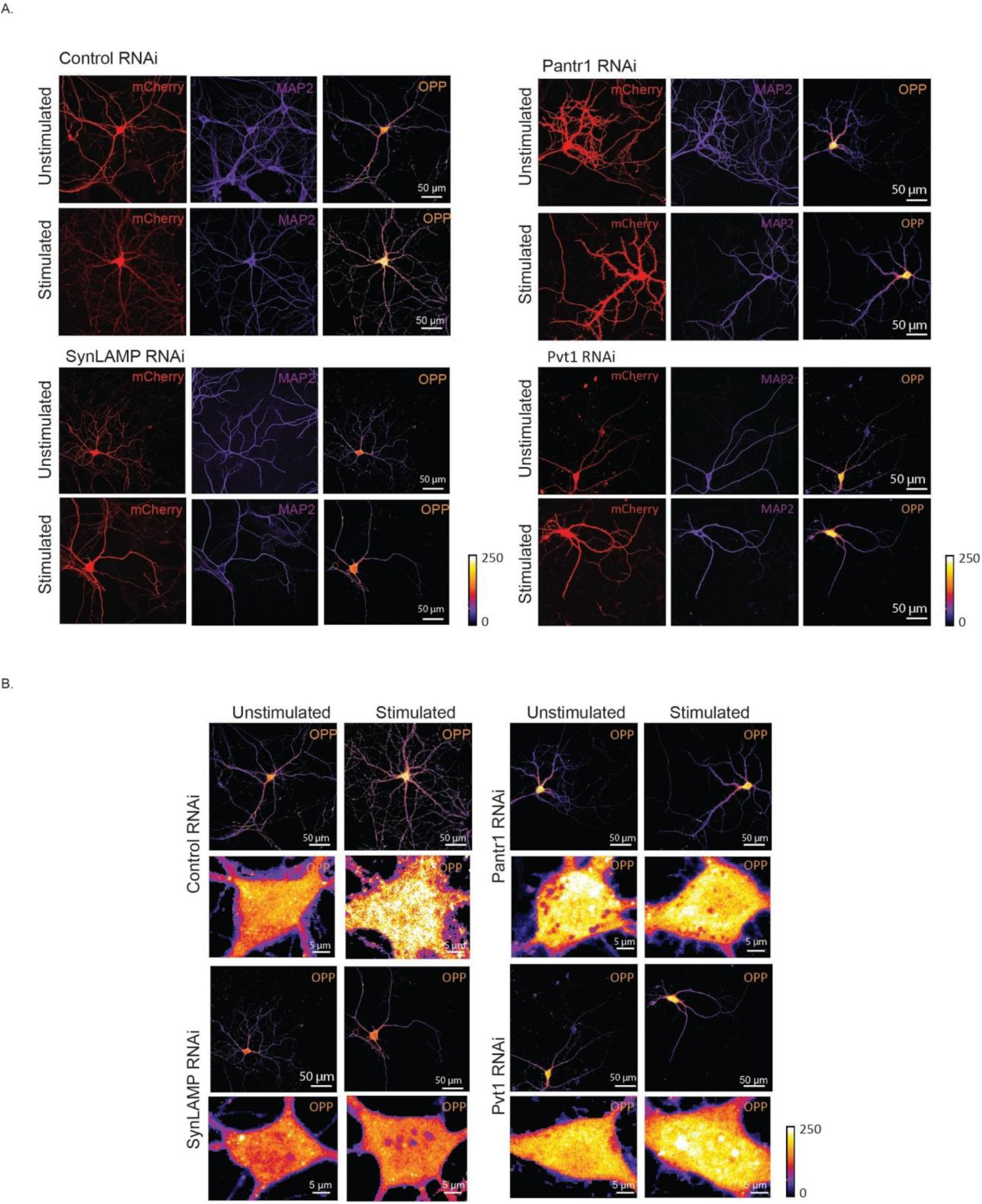
Activity-regulated protein synthesis in hippocampal neurons following knockdown of Pantr1, SynLAMP and Pvt1. (**A**) Cultured hippocampal neurons (DIV 10) were transduced with lentivirus expressing shRNAs as indicated. Neurons (DIV 21 - 23) were stimulated with Glutamate in presence of Actinomycin D. Protein synthesis from the mCherry (red) expressing transduced neurons was assessed by Puromycin incorporation (O-propargyl-puromycin or OPP; an alkyne analog of puromycin) and dendrites (blue) were visualized by MAP2 immunostaining. Intensity of fluorescence observed in neurons is a direct readout of OPP incorporation due to protein synthesis. Differential pixel intensity has been represented as a heat map (low: blue to high: white). Calibration bar and scale bars as indicated. (**B**) Images showing the extent of OPP incorporation in the soma of neurons expressing shRNA as indicated. Differential pixel intensity has been represented as a heat map. Calibration bar and scale bars as indicated. Also see Figure 5.

**Figure S8:**
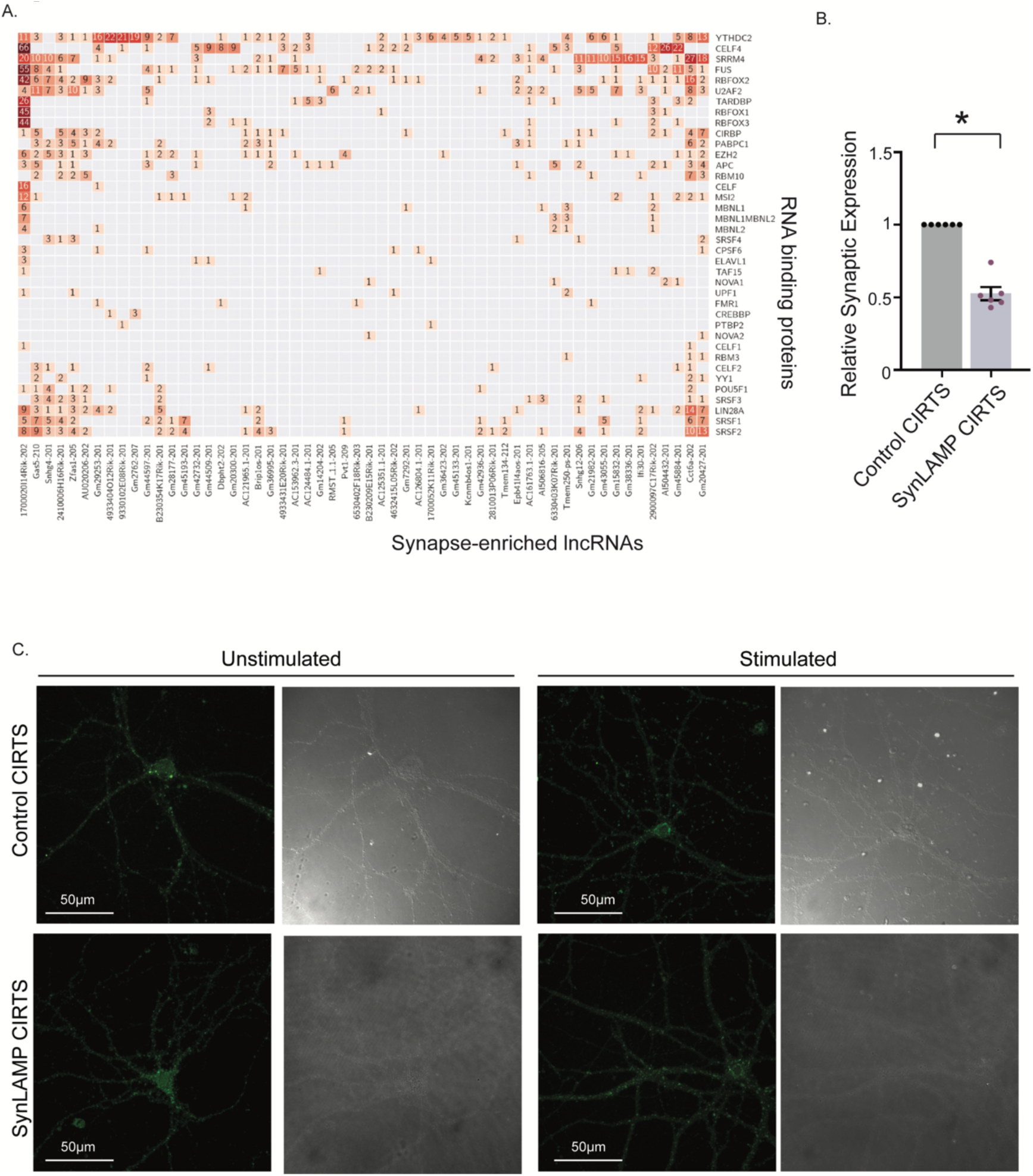
Synapse-enriched lncRNA-RBP interaction following contextual fear conditioning. **(A)** RBP binding motif present in indicated synapse-enriched lncRNAs as detected by *in silico* analysis using the POSTAR3 database. **(B)** Relative expression of SynLAMP in the synaptoneurosomes from the hippocampus of UC-CIRTS-injected, and SynLAMP-CIRTS-injected adult mice, post contextual fear conditioning.**(C)** Representative photomicrographs of Puro-PLA signal in cultured hippocampal neurons expressing Control-CIRTS and SynLAMP-CIRTS. Scale as indicated. Also see Figure 6 and Figure 7.

## References

Ashraf, S.I., McLoon, A.L., Sclarsic, S.M., and Kunes, S. (2006). Synaptic protein synthesis associated with memory is regulated by the RISC pathway in Drosophila. Cell 124, 191–205.

Askarian-Amiri, M.E., Crawford, J., French, J.D., Smart, C.E., Smith, M.A., Clark, M.B., Ru, K., Mercer, T.R., Thompson, E.R., Lakhani, S.R., et al. (2011). SNORD-host RNA Zfas1 is a regulator of mammary development and a potential marker for breast cancer. RNA 17, 878– 891.

Banerjee, S., Neveu, P., and Kosik, K.S. (2009). A coordinated local translational control point at the synapse involving relief from silencing and MOV10 degradation. Neuron 64, 871–884.

Belly, A., Moreau-Gachelin, F., Sadoul, R., and Goldberg, Y. (2005). Delocalization of the multifunctional RNA splicing factor TLS/FUS in hippocampal neurones: exclusion from the nucleus and accumulation in dendritic granules and spine heads. Neurosci. Lett. 379, 152– 157.

Bernard, D., Prasanth, K. V., Tripathi, V., Colasse, S., Nakamura, T., Xuan, Z., Zhang, M.Q., Sedel, F., Jourdren, L., Coulpier, F., et al. (2010). A long nuclear-retained non-coding RNA regulates synaptogenesis by modulating gene expression. EMBO J. 29, 3082–3093.

Bourtchouladze, R., Abel, T., Berman, N., Gordon, R., Lapidus, K., and Kandel, E.R. (1998). Different Training Procedures Recruit Either One or Two Critical Periods for Contextual Memory Consolidation, Each of Which Requires Protein Synthesis and PKA. Learn. Mem. 5, 365–374.

Bramham, C.R., and Wells, D.G. (2007). Dendritic mRNA: transport, translation and function. Nat. Rev. Neurosci. 8, 776–789.

Briggs, J.A., Wolvetang, E.J., Mattick, J.S., Rinn, J.L., and Barry, G. (2015). Mechanisms of Long Non-coding RNAs in Mammalian Nervous System Development, Plasticity, Disease, and Evolution. Neuron 88, 861–877.

Bu, D., Yu, K., Sun, S., Xie, C., Skogerbø, G., Miao, R., Xiao, H., Liao, Q., Luo, H., Zhao, G., et al. (2012). NONCODE v3.0: Integrative annotation of long noncoding RNAs. Nucleic Acids Res. 40, 210–215.

Cajigas, I.J., Tushev, G., Will, T.J., Tom Dieck, S., Fuerst, N., and Schuman, E.M. (2012). The Local Transcriptome in the Synaptic Neuropil Revealed by Deep Sequencing and High-Resolution Imaging. Neuron 74, 453–466.

Castejón, O.J., Fuller, L., and Dailey, M.E. (2004). Localization of synapsin-I and PSD-95 in developing postnatal rat cerebellar cortex. Dev. Brain Res. 151, 25–32.

Chen, N., Zhang, Y., Adel, M., Kuklin, E.A., Reed, M.L., Mardovin, J.D., Bakthavachalu, B., VijayRaghavan, K., Ramaswami, M., and Griffith, L.C. (2022). Local translation provides the asymmetric distribution of CaMKII required for associative memory formation. Curr. Biol. 32, 2730–2738.e5.

Clemson, C.M., Hutchinson, J.N., Sara, S.A., Ensminger, A.W., Fox, A.H., Chess, A., and Lawrence, J.B. (2009). An Architectural Role for a Nuclear Noncoding RNA: NEAT1 RNA Is Essential for the Structure of Paraspeckles. Mol. Cell 33, 717–726.

Conde-Dusman, M.J., Dey, P.N., Elía-Zudaire, Ó., Rabaneda, L.G., García-Lira, C., Grand, T., Briz, V., Velasco, E.R., Andero, R., Niñerola, S., et al. (2021). Control of protein synthesis and memory by glun3a-nmda receptors through inhibition of git1/mtorc1 assembly. Elife 10.

Darnell, R.B. (2013). RNA Protein Interaction in Neurons. 10.1146/Annurev-Neuro-062912-114322 36, 243–270.

Dieck, S.T., Muller, A., Nehring, A., Hinz, F.I., Bartnik, I., Schuman, E.M., and Dieterich, D.C. (2012). Metabolic Labeling with Noncanonical Amino Acids and Visualization by Chemoselective Fluorescent Tagging. Curr. Protoc. Cell Biol. 56, 7.11.1–7.11.29.

Dong, X.Y., Guo, P., Boyd, J., Sun, X., Li, Q., Zhou, W., and Dong, J.T. (2009). Implication of snoRNA U50 in human breast cancer. J. Genet. Genomics 36, 447–454.

Dunham, I., Kundaje, A., Aldred, S.F., Collins, P.J., Davis, C.A., Doyle, F., Epstein, C.B., Frietze, S., Harrow, J., Kaul, R., et al. (2012). An Integrated Encyclopedia of DNA Elements in the Human Genome. Nature 489, 57.

Espadas, I., Wingfield, J.L., Nakahata, Y., Chanda, K., Grinman, E., Ghosh, I., Bauer, K.E., Raveendra, B., Kiebler, M.A., Yasuda, R., et al. (2024). Synaptically-targeted long non-coding RNA SLAMR promotes structural plasticity by increasing translation and CaMKII activity. Nat. Commun. 15.

Fanselow, M.S. (2000). Contextual fear, gestalt memories, and the hippocampus. Behav. Brain Res. 110, 73–81.

Fernandez-Moya, S.M., Bauer, K.E., and Kiebler, M.A. (2014). Meet the players: Local translation at the synapse. Front. Mol. Neurosci. 7, 84.

Fletcher, T.L., De Camilli, P., and Banker, G. (1994). Synaptogenesis in hippocampal cultures: Evidence indicating that axons and dendrites become competent to form synapses at different stages of neuronal development. J. Neurosci. 14, 6695–6706.

Frey, U., and Morris, R.G. (1997). Synaptic tagging and long-term potentiation. Nature 385, 533–536.

Goff, L.A., Groff, A.F., Sauvageau, M., Trayes-Gibson, Z., Sanchez-Gomez, D.B., Morse, M., Martin, R.D., Elcavage, L.E., Liapis, S.C., Gonzalez-Celeiro, M., et al. (2015). Spatiotemporal expression and transcriptional perturbations by long noncoding RNAs in the mouse brain. Proc. Natl. Acad. Sci. U. S. A. 112, 6855–6862.

Grinman, E., Nakahata, Y., Avchalumov, Y., Espadas, I., Swarnkar, S., Yasuda, R., and Puthanveettil, S. V (2021). Activity-regulated synaptic targeting of lncRNA ADEPTR mediates structural plasticity by localizing Sptn1 and AnkB in dendrites. Sci Adv 7, 605–621.

Hafezqorani, S., Houdjedj, A., Arici, M., Said, A., and Kazan, H. (2019). RBPSponge: genome-wide identification of lncRNAs that sponge RBPs. Bioinformatics 35, 4760–4763.

Hall, B.J., and Ghosh, A. (2008). Regulation of AMPA receptor recruitment at developing synapses. Trends Neurosci. 31, 82–89.

Harrow, J., Frankish, A., Gonzalez, J.M., Tapanari, E., Diekhans, M., Kokocinski, F., Aken, B.L., Barrell, D., Zadissa, A., Searle, S., et al. (2012). GENCODE: The reference human genome annotation for the ENCODE project. Genome Res. 22, 1760–1774.

Heraud-Farlow, J.E., Sharangdhar, T., Li, X., Pfeifer, P., Tauber, S., Orozco, D., Hörmann, A., Thomas, S., Bakosova, A., Farlow, A.R., et al. (2013). Staufen2 regulates neuronal target RNAs. Cell Rep. 5, 1511–1518.

Ho, V.M., Lee, J.A., and Martin, K.C. (2011). The Cell Biology of Synaptic Plasticity. Science (80-. ). 334, 623–628.

Holt, C.E., and Schuman, E.M. (2013). The Central Dogma Decentralized: New Perspectives on RNA Function and Local Translation in Neurons. Neuron 80, 648–657.

Hotulainen, P., and Hoogenraad, C.C. (2010). Actin in dendritic spines: connecting dynamics to function. J. Cell Biol. 189, 619–629.

Kandel, E.R. (2001). The Molecular Biology of Memory Storage: A Dialogue Between Genes and Synapses. Science. 294, 1030–1038.

Keihani, S., Kluever, V., Mandad, S., Bansal, V., Rahman, R., Fritsch, E., Caldi Gomes, L., Gärtner, A., Kügler, S., Urlaub, H., et al. (2019). The long noncoding RNA neuroLNC regulates presynaptic activity by interacting with the neurodegeneration-associated protein TDP-43. Sci. Adv. 5 (12), eaay2670.

Kennedy, M.B., Beale, H.C., Carlisle, H.J., and Washburn, L.R. (2005). Integration of biochemical signalling in spines. Nat. Rev. Neurosci. 6, 423–434.

Khalil, A.M., Guttman, M., Huarte, M., Garber, M., Raj, A., Morales, D.R., Thomas, K., Presser, A., Bernstein, B.E., Van Oudenaarden, A., et al. (2009). Many human large intergenic noncoding RNAs associate with chromatin-modifying complexes and affect gene expression. Proc. Natl. Acad. Sci. U. S. A. 106, 11667–11672.

Kiebler, M.A., and Bassell, G.J. (2006). Neuronal RNA Granules: Movers and Makers. Neuron 51, 685–690.

Klann, E., and Dever, T.E. (2004). Biochemical mechanisms for translational regulation in synaptic plasticity. Nat. Rev. Neurosci. 5, 931–942.

Krichevsky, A.M., and Kosik, K.S. (2001). Neuronal RNA granules: A link between RNA localization and stimulation-dependent translation. Neuron 32, 683–696.

Kye, M.J., Liu, T., Levy, S.F., Nan, L.X., Groves, B.B., Bonneau, R., Lao, K., and Kosik, K.S. (2007). Somatodendritic microRNAs identified by laser capture and multiplex RT-PCR. RNA 13, 1224–1234.

De Leeuw, F., Zhang, T., Wauquier, C., Huez, G., Kruys, V., and Gueydan, C. (2007). The cold-inducible RNA-binding protein migrates from the nucleus to cytoplasmic stress granules by a methylation-dependent mechanism and acts as a translational repressor. Exp. Cell Res. 313, 4130–4144.

Li, D., Zhang, J., Wang, M., Li, X., Gong, H., Tang, H., Chen, L., Wan, L., and Liu, Q. (2018). Activity dependent LoNA regulates translation by coordinating rRNA transcription and methylation. Nat. Commun. 9(1):1726.doi:10.1038/s41467-018-04072-4

Liau, W.S., Zhao, Q., Bademosi, A., Gormal, R.S., Gong, H., Marshall, P.R., Periyakaruppiah, A., Madugalle, S.U., Zajaczkowski, E.L., Leighton, L.J., et al. (2023). Fear extinction is regulated by the activity of long noncoding RNAs at the synapse. Nat. Commun. 2023 141 14, 1–16.

Logan, S.M., and Storey, K.B. (2020). Cold-inducible RNA-binding protein Cirp, but not Rbm3, may regulate transcript processing and protection in tissues of the hibernating ground squirrel. Cell Stress Chaperones 25, 857–868.

Luo, Y., Hitz, B.C., Gabdank, I., Hilton, J.A., Kagda, M.S., Lam, B., Myers, Z., Sud, P., Jou, J., Lin, K., et al. (2020). New developments on the Encyclopedia of DNA Elements (ENCODE) data portal. Nucleic Acids Res. 48, D882–D889.

Martin, K.C., and Ephrussi, A. (2009). mRNA Localization: Gene Expression in the Spatial Dimension. Cell 136, 719–730.

Matsuzaki, M., Honkura, N., Ellis-Davies, G.C.R., and Kasai, H. (2004). Structural basis of long-term potentiation in single dendritic spines. Nature 429, 761–766.

Mattick, J.S. (2023). RNA out of the mist. Trends Genet. 39, 187–207.

Mercer, T.R., Dinger, M.E., Mariani, J., Kosik, K.S., Mehler, M.F., and Mattick, J.S. (2007). Noncoding RNAs in Long-Term Memory Formation. The Neuroscientist 14, 434–445

Mercer, T.R., Dinger, M.E., Sunkin, S.M., Mehler, M.F., and Mattick, J.S. (2008). Specific expression of long noncoding RNAs in the mouse brain. Proc. Natl. Acad. Sci. 105, 716–721.

Mercer, T.R., Dinger, M.E., and Mattick, J.S. (2009). Long non-coding RNAs: insights into functions. Nat Rev Genet 10, 155–159.

Mercer, T.R., Gerhardt, D.J., Dinger, M.E., Crawford, J., Trapnell, C., Jeddeloh, J.A., Mattick, J.S., and Rinn, J.L. (2012). Targeted RNA sequencing reveals the deep complexity of the human transcriptome. Nat. Biotechnol. 30, 99.

Miller, S., Yasuda, M., Coats, J.K., Jones, Y., Martone, M.E., and Mayford, M. (2002). Disruption of Dendritic Translation of CaMKIIα Impairs Stabilization of Synaptic Plasticity and Memory Consolidation. Neuron 36, 507–519.

Mondal, T., Rasmussen, M., Pandey, G.K., Isaksson, A., and Kanduri, C. (2010). Characterization of the RNA content of chromatin. Genome Res. 20, 899–907.

Montarolo, P.G. (1986). A critical period for macromolecular synthesis in long-term heterosynaptic facilitation in Aplysia. Science (80-. ). 234, 1249–1254.

Mourtada-Maarabouni, M., Pickard, M.R., Hedge, V.L., Farzaneh, F., and Williams, G.T. (2008). GAS5, a non-protein-coding RNA, controls apoptosis and is downregulated in breast cancer. Oncogene 2009 282 28, 195–208.

Mukherjee, N., Calviello, L., Hirsekorn, A., De Pretis, S., Pelizzola, M., and Ohler, U. (2016). Integrative classification of human coding and noncoding genes through RNA metabolism profiles. Nat. Struct. Mol. Biol. 2016 241 24, 86–96.

Nakayama, A.Y., and Luo, L. (2000). Intracellular signalling pathways that regulate dendritic spine morphogenesis. Hippocampus 10, 582–586.

O’Brien, R.J., Kamboj, S., Ehlers, M.D., Rosen, K.R., Fischbach, G.D., and Huganir, R.L. (1998). Activity-dependent modulation of synaptic AMPA receptor accumulation. Neuron 21, 1067–1078.

O’Donnell, C., and Sejnowski, T.J. (2014). Selective Memory Generalization by Spatial Patterning of Protein Synthesis. Neuron 82, 398–412.

Paradis, S., Harrar, D.B., Lin, Y., Koon, A.C., Hauser, J.L., Griffith, E.C., Zhu, L., Brass, L.F., Chen, C., and Greenberg, M.E. (2007). An RNAi-Based Approach Identifies Molecules Required for Glutamatergic and GABAergic Synapse Development. Neuron 53, 217–232.

Pelczar, P., and Filipowicz, W. (1998). The Host Gene for Intronic U17 Small Nucleolar RNAs in Mammals Has No Protein-Coding Potential and Is a Member of the 5′-Terminal Oligopyrimidine Gene Family. Mol. Cell. Biol. 18, 4509–4518.

Phillips, R.G., and LeDoux, J.E. (1992). Differential Contribution of Amygdala and Hippocampus to Cued and Contextual Fear Conditioning. Behav. Neurosci. 106, 274–285.

Piwecka, M., Glažar, P., Hernandez-Miranda, L.R., Memczak, S., Wolf, S.A., Rybak-Wolf, A., Filipchyk, A., Klironomos, F., Jara, C.A.C., Fenske, P., et al. (2017). Loss of a mammalian circular RNA locus causes miRNA deregulation and affects brain function. Science (80-. ). 357.

Rao, A., and Steward, O. (1991). Evidence that Protein Constituents of Postsynaptic Membrane Specializations Are Locally Synthesized: Analysis of Proteins Synthesized within Synaptosomes. J. Neurosci. 1, 2881–2895.

Ray, D., Kazan, H., Cook, K.B., Weirauch, M.T., Najafabadi, H.S., Li, X., Gueroussov, S., Albu, M., Zheng, H., Yang, A., et al. (2013). A compendium of RNA-binding motifs for decoding gene regulation. Nat. 2013 4997457 499, 172–177.

Rybak-Wolf, A., Stottmeister, C., Glažar, P., Jens, M., Pino, N., Giusti, S., Hanan, M., Behm, M., Bartok, O., Ashwal-Fluss, R., et al. (2015). Circular RNAs in the Mammalian Brain Are Highly Abundant, Conserved, and Dynamically Expressed. Mol. Cell 58, 870–885.

Sahadevan, S., Hembach, K.M., Tantardini, E., Pérez-Berlanga, M., Hruska-Plochan, M., Megat, S., Weber, J., Schwarz, P., Dupuis, L., Robinson, M.D., et al. (2021). Synaptic FUS accumulation triggers early misregulation of synaptic RNAs in a mouse model of ALS. Nat. Commun. 2021 121 12, 1–17.

Samaddar, S., and Banerjee, S. (2021). Far from the nuclear crowd: Cytoplasmic lncRNA and their implications in synaptic plasticity and memory. Neurobiol. Learn. Mem. 185, 107522.

Sambandan, S., Akbalik, G., Kochen, L., Rinne, J., Kahlstatt, J., Glock, C., Tushev, G., Alvarez-Castelao, B., Heckel, A., and Schuman, E.M. (2017). Activity-dependent spatially localized miRNA maturation in neuronal dendrites. Science (80-. ). 355, 634 LP – 637.

Schieweck, R., Ninkovic, J., and Kiebler, M.A. (2020). RNA-binding proteins balance brain function in health and disease. Physiol. Rev.

Schratt, G.M., Nigh, E.A., Chen, W.G., Hu, L., and Greenberg, M.E. (2004). BDNF Regulates the Translation of a Select Group of mRNAs by a Mammalian Target of Rapamycin-Phosphatidylinositol 3-Kinase-Dependent Pathway during Neuronal Development. J. Neurosci. 24, 7366–7377.

Schratt, G.M., Tuebing Fabian, Nigh, E.A., Kane, C.G., Sabatini, M.E., Kiebler, M., and Greenberg, M.E. (2006). A brain-specific microRNA regulates dendritic spine development. Nature 439, 283–289.

Schubert, V., and Dotti, C.G. (2007). Transmitting on actin: synaptic control of dendritic architecture. J. Cell Sci. 120, 205–212.

Shiina, N., Shinkura, K., and Tokunaga, M. (2005). A novel RNA-binding protein in neuronal RNA granules: Regulatory machinery for local translation. J. Neurosci. 25, 4420–4434.

Siegel, G., Obernosterer, G., Fiore, R., Oehmen, M., Bicker, S., Christensen, M., Khudayberdiev, S., Leuschner, P.F., Busch, C.J.L., Kane, C., et al. (2009). A functional screen implicates microRNA-138-dependent regulation of the depalmitoylation enzyme APT1 in dendritic spine morphogenesis. Nat. Cell Biol. 11, 705–716.

Sévigny, M., Julien, I.B., Venkatasubramani, J.P., Hui, J.B., Dutchak, P.A., and Sephton, C.F. (2020). FUS contributes to mTOR-dependent inhibition of translation. J. Biol. Chem. 295, 18459–18473.

Srinivasan, B., Samaddar, S., Mylavarapu, S.V.S., Clement, J.P., and Banerjee, S. (2021). Homeostatic scaling is driven by a translation-dependent degradation axis that recruits miRISC remodeling. PLoS Biol. 19 (11),e3001432.

Statello, L., Guo, C.J., Chen, L.L., and Huarte, M. (2021). Gene regulation by long non-coding RNAs and its biological functions. Nat. Rev. Mol. Cell Biol. 22, 96–118.

Sutton, M.A., and Schuman, E.M. (2006). Dendritic Protein Synthesis, Synaptic Plasticity, and Memory. Cell 127, 49–58.

Swarnkar, S., Avchalumov, Y., Espadas, I., Grinman, E., Liu, X. an, Raveendra, B.L., Zucca, A., Mediouni, S., Sadhu, A., Valente, S., et al. (2021). Molecular motor protein KIF5C mediates structural plasticity and long-term memory by constraining local translation. Cell Rep. 36, 109369.

Threadgill, R., Bobb, K., and Ghosh, A. (1997). Regulation of Dendritic Growth and Remodeling by Rho, Rac, and Cdc42. Neuron 19, 625–634.

Tolias, K.F., Duman, J.G., and Um, K. (2011). Control of synapse development and plasticity by Rho GTPase regulatory proteins. Prog. Neurobiol. 94, 133–148.

Tom Dieck, S., Kochen, L., Hanus, C., Heumüller, M., Bartnik, I., Nassim-Assir, B., Merk, K., Mosler, T., Garg, S., Bunse, S., et al. (2015). Direct visualization of newly synthesized target proteins in situ. Nat. Methods 12, 411–414.

Tong, G., Endersfelder, S., Rosenthal, L.M., Wollersheim, S., Sauer, I.M., Bührer, C., Berger, F., and Schmitt, K.R.L. (2013). Effects of moderate and deep hypothermia on RNA-binding proteins RBM3 and CIRP expressions in murine hippocampal brain slices. Brain Res. 1504, 74–84.

Tsai, M.C., Manor, O., Wan, Y., Mosammaparast, N., Wang, J.K., Lan, F., Shi, Y., Segal, E., and Chang, H.Y. (2010). Long noncoding RNA as modular scaffold of histone modification complexes. Science (80-. ). 329, 689–693.

Tushev, G., Glock, C., Heumüller, M., Biever, A., Jovanovic, M., and Schuman, E.M. (2018). Alternative 3’ UTRs Modify the Localization, Regulatory Potential, Stability, and Plasticity of mRNAs in Neuronal Compartments. Neuron 98, 495–511.e6.

Udagawa, T., Fujioka, Y., Tanaka, M., Honda, D., Yokoi, S., Riku, Y., Ibi, D., Nagai, T., Yamada, K., Watanabe, H., et al. (2015). FUS regulates AMPA receptor function and FTLD/ALS-associated behaviour via GluA1 mRNA stabilization. Nat. Commun. 2015 61 6, 1–13.

Wang, D.O., Kim, S.M., Zhao, Y., Hwang, H., Miura, S.K., Sossin, W.S., and Martin, K.C. (2009). Synapse- and stimulus-specific local translation during long-term neuronal plasticity. Science (80-. ). 324, 1536–1540.

Wei, W., Zhao, Q., Wang, Z., Liau, W.S., Basic, D., Ren, H., Marshall, P.R., Zajaczkowski, E.L., Leighton, L.J., Madugalle, S.U., et al. (2022). ADRAM is an experience-dependent long noncoding RNA that drives fear extinction through a direct interaction with the chaperone protein 14-3-3. Cell Rep. 38, 110546.

Wu, G.Y., Malinow, R., and Cline, H.T. (1996). Maturation of a Central Glutamatergic Synapse. Science 274, 972–976.

Xia, Z., Zheng, X., Zheng, H., Liu, X., Yang, Z., and Wang, X. (2012). Cold-inducible RNA-binding protein (CIRP) regulates target mRNA stabilization in the mouse testis. FEBS Lett. 586, 3299–3308.

Xie, C., Yuan, J., Li, H., Li, M., Zhao, G., Bu, D., Zhu, W., Wu, W., Chen, R., and Zhao, Y. (2014). NONCODEv4: Exploring the world of long non-coding RNA genes. Nucleic Acids Res. 42, D98–D103.

Yasuda, K., Zhang, H., Loiselle, D., Haystead, T., Macara, I.G., and Mili, S. (2013). The RNA-binding protein Fus directs translation of localized mRNAs in APC-RNP granules. J. Cell Biol. 203, 737–746.

You, X., Vlatkovic, I., Babic, A., Will, T., Epstein, I., Tushev, G., Akbalik, G., Wang, M., Glock, C., Quedenau, C., et al. (2015). Neural circular RNAs are derived from synaptic genes and regulated by development and plasticity. Nat. Neurosci. 2015 184 18, 603–610.

Zhao, W., Zhang, S., Zhu, Y., Xi, X., Bao, P., Ma, Z., Kapral, T.H., Chen, S., Zagrovic, B., Yang, Y.T., et al. (2021). POSTAR3: an updated platform for exploring post-transcriptional regulation coordinated by RNA-binding proteins. Nucleic Acids Res. 50.

Zhao, Y., Liu, S., Zhou, L., Li, X., Meng, Y., Li, Y., Li, L., Jiao, B., Bai, L., Yu, Y., et al. (2019). Aberrant shuttling of long noncoding RNAs during the mitochondria-nuclear crosstalk in hepatocellular carcinoma cells. Am. J. Cancer Res. 9, 999.

Zhou, Y., Lu, H., Liu, Y., Zhao, Z., Zhang, Q., Xue, C., Zou, Y., Cao, Z., and Luo, W. (2021). Cirbp-PSD95 axis protects against hypobaric hypoxia-induced aberrant morphology of hippocampal dendritic spines and cognitive deficits. Mol. Brain 14, 1–17.

Zhu, Y., Xu, G., Yang, Y.T., Xu, Z., Chen, X., Shi, B., Xie, D., Lu, Z.J., and Wang, P. (2019). POSTAR2: Deciphering the post-Transcriptional regulatory logics. Nucleic Acids Res. 47, D203–D211.

Zito, K., Knott, G., Shepherd, G.M.G., Shenolikar, S., and Svoboda, K. (2004). Induction of Spine Growth and Synapse Formation by Regulation of the Spine Actin Cytoskeleton. Neuron 44, 321–334

